# Quantifying Population Reversibility of Sensor Performance in Multi-Cycle Single-Sensor Recovery Assay

**DOI:** 10.64898/2025.12.25.696491

**Authors:** Geffen Rosenberg, Gili Bisker

**Affiliations:** School of Biomedical Engineering, Faculty of Engineering, Tel Aviv University, Tel Aviv 6997801, Israel; Center for Physics and Chemistry of Living Systems, Tel Aviv University, Tel Aviv 6997801, Israel; Center for Nanoscience and Nanotechnology, Tel Aviv University, Tel Aviv 6997801, Israel; Center for Light-Matter Interaction, Tel Aviv University, Tel Aviv 6997801, Israel; Sagol School of Neuroscience, Tel Aviv University, Tel Aviv 6997801, Israel

**Keywords:** single-walled carbon nanotubes, fluorescence sensors, near-infrared, sensor reversibility, sensor recovery, spatiotemporal sensors

## Abstract

Quantitative chemical imaging requires sensors that reliably recover across repeated exposures. While solution-phase bulk measurements provide an averaged response of the sensor population, imaging at the single-sensor level enables mapping of biological processes with spatiotemporal resolution, revealing localized events and interaction sites. To translate such imaging into calibrated measurements, sensor variability under repeated analyte exposures must be analyzed. This work introduces a generic workflow that combines an automated microfluidic flow imaging platform with systematic characterization of the response, recovery, and reversibility of individual nanosensors across multi-cycle challenges. For representative implementation, three near-infrared fluorescent single-walled carbon nanotube (SWCNT) sensor models are tested, each with a distinct functionalization rendering it optically responsive to a corresponding exemplar target: dopamine, thiocholine, or serotonin. While first-cycle responses averaged over the entire field of view recapitulate ensemble calibration, single-sensor analysis uncovers broad heterogeneity in response magnitude, signal recovery, and reversibility across hundreds of individual SWCNTs under repeated exposure and wash cycles. To compare performance across cycles, a standardized Population Reversibility Score based on Kullback-Leibler Divergence is introduced, condensing response distributions into a single cycle- and concentration-dependent, quantitative metric. This framework is generally applicable to other sensor-analyte systems with transient readouts, guiding optimization for spatiotemporal analyte mapping.

## 1. Introduction

Monitoring chemical messengers in living systems requires sensors that are not only sensitive and selective but also capable of providing spatiotemporal information on the presence of these messenger biomolecules.^[1,2]^ From synaptic neurotransmission to paracrine signaling and oxidative bursts, these events are transient, localized, and repeated, favoring imaging-based readouts with cellular and subcellular resolution over conventional ensemble assays.^[3–5]^ In such cases, sensors must report reliably across many exposure-wash cycles, so that quantitative maps of concentration and dynamics can be trusted over time.^[6,7]^ Given single-sensor heterogeneity and incomplete recovery after repeated challenges, there is a need for standardized metrics to compare reversibility across constructs, doses, and experimental conditions.

Within this context, single-walled carbon nanotubes (SWCNTs) have emerged as a powerful optical sensing platform, capable of detecting and continuously monitoring a variety of target analytes. Structurally, a SWCNT can be considered a rolled-up single-layer graphene sheet, with a diameter of approximately 1-2 nm and varying lengths reaching up to hundreds of micrometers.^[8–10]^ The roll-up vector of the graphene sheet defines the SWCNT’s chirality, which in turn determines the SWCNT’s electronic band structure and thus its optical and fluorescence properties.^[8,9,11]^ Semiconducting SWCNTs fluoresce in the near-infrared (NIR) range, between 900 nm and 1400 nm,^[12,13]^ which coincides with the optical transparency window for biological samples. This overlap enables high signal-to-noise ratio imaging, as it benefits from reduced background autofluorescence, absorption, and scattering.^[14–16]^ Combined with their stable fluorescence, which does not suffer from photobleaching or blinking,^[17,18]^ and their biocompatibility when appropriately functionalized,^[19–24]^ SWCNTs are highly attractive as optical nanosensors.^[17,25–28]^ Analyte recognition and signal transduction arise from the surface functionalization of the SWCNTs.^[29–31]^ By suspending pristine or defect-induced^[32–37]^ SWCNTs with different polymers,^[38–41]^ peptides,^[42–45]^ or DNA,^[46–49]^ the corona chemistry mediates specific interactions with target analytes, leading to a detectable modulation of the emitted SWCNT fluorescence.^[21,50,51]^ This approach has yielded SWCNT sensors for a wide variety of analytes, including proteins,^[38,39,52–55]^ enzymes,^[41,46,56–59]^ neurotransmitters,^[47,48,60–65]^ and hormones.^[40,66,67]^

Traditionally, SWCNT sensing studies quantify ensemble fluorescence responses in aqueous solutions, averaging over large, heterogeneous sensor populations.^[68]^ However, the nanoscale dimensions of SWCNT also enable single-sensor imaging with exceptional spatiotemporal resolution for detecting analyte presence. Recent work has leveraged this capability to monitor plant health,^[69–71]^ track reactive oxygen species in photoaged skin cells,^[72]^ visualize neurotransmitter release in neuronal systems,^[62,66,73–80]^ and probe the gastrointestinal tract within *C. elegans* worms.^[81]^ For example, Lee *et al.* embedded dopamine (DA) sensors around differentiating human induced pluripotent stem cells (hiPSC)-derived dopaminergic neurons and, with ∼200 ms temporal and micrometer spatial resolution, mapped release “hotspots”, revealing the differences in DA release between healthy and GBA1-mutant neurons across differentiation stages.^[77]^ Mun *et al.* developed an oxytocin sensor and imaged stimulus-evoked release in acute brain slices with ∼250 ms resolution, enabling regional comparisons of hotspot number and amplitude.^[66]^ Moreover, Dinarvand *et al.* demonstrated SWCNT-based serotonin sensors that enabled real-time, sensitive spatiotemporal imaging of serotonin release from individual human platelets, revealing localized secretion hotspots and dynamic release kinetics at subcellular resolution.^[62]^ In practice, the same SWCNT endures multiple analyte exposures, so quantitative readouts require reversibility, recovery, and consistent response. While reversibility has been shown qualitatively for several SWCNT sensor constructs, a quantitative analysis of single-sensor recovery across repeated challenges is lacking. Moreover, without explicit characterization of single-sensor performance heterogeneity, ensemble-based calibration and measurement can misrepresent local sensing behavior during high-resolution imaging.

Here, we address these gaps by systematically characterizing response distributions and multi-cycle reversibility of individual SWCNTs in a controlled microfluidic platform. We study three known single-strand DNA (ssDNA)-functionalized SWCNT-analyte pairs, targeting dopamine (DA),^[47]^ thiocholine (TCh),^[46]^ and serotonin (5-HT).^[48]^ Using automated flow control and NIR imaging, we compare first-cycle immobilized sensor responses to bulk calibrations, investigate sensor-to-sensor heterogeneity across hundreds of regions of interest, and track recovery over repeated exposure-wash cycles across a wide range of analyte concentrations within the sensors’ dynamic range. To standardize comparisons across cycles and concentrations, we introduce a cycle- and concentration-dependent Population Reversibility Score based on the Kullback–Leibler divergence, also known as relative entropy, which quantifies the distinguishability between two probability distributions. In our context, we measure the deviation of each cycle’s response from its first-cycle reference, thus condensing the statistical differences in response distributions of heterogeneous SWCNT populations into a single metric of reversibility. Our Reversibility Score elucidates the effects of analyte polymerization, incomplete washout, and potential corona modifications, thereby establishing design and analysis principles for reliable, quantitative single-sensor high-resolution imaging with nanosensors.

## 2. Results

### 2.1 Initial bulk characterization of SWCNT sensor response

HipCO SWCNTs were suspended with single-strand DNA (ssDNA) corona, DNA1, DNA2, and DNA3 (see the full sequences in the Experimental Section), to render them optically responsive to Dopamine (DA)^[47]^, thiocholine (TCh),^[46]^ and serotonin (5-HT),^[48]^ respectively. The DNA1-SWCNT sensor for DA was discovered via corona phase molecular recognition screening of polymer-wrapped SWCNTs, where DNA1 ssDNA emerged as a lead corona that yields a significant turn-on fluorescence response to dopamine with high sensitivity, attributed to dopamine-induced conformational changes in the DNA corona that increase the SWCNT’s fluorescence quantum yield.^[47]^ The DNA2-SWCNT sensor for TCh was identified via a DNA-SWCNT library screen for cholinesterase (ChE) activity, in which DNA2-SWCNTs showed a strong, selective fluorescence increase in response to thiocholine, the enzymatic hydrolysis product of acetylthiocholine, with negligible responses to the enzyme or substrate alone, establishing DNA2-SWCNTs as a reporter of ChE activity and inhibition.^[46]^ DNA3-SWCNT was found through machine learning models trained on spectral data sets of numerous DNA-SWCNT conjugates to predict high-response candidates, which yielded several ssDNA-SWCNTs, including DNA3, predicted to exhibit a strong fluorescence response to 5-HT.^[48]^ In experimental validation, DNA3-SWCNT showed the strongest response to 5-HT and was therefore selected for our study.

Successful functionalization was confirmed by the distinguishable peaks of the different SWCNTs chiralities in the absorption spectra (**Figure S1**) and in the excitation-emission maps of their NIR-fluorescence (**Figure S2**). Subsequently, the fluorescence responses of the suspensions to different concentrations of the corresponding analytes were recorded (**Figure S3**), showing an increase in fluorescence intensity with increasing analyte concentrations for all analytes tested. For DNA1-SWCNT, the response reached saturation at ∼50 µM DA, but declined above 100 µM DA, likely due to DA polymerization^[47,82]^. The normalized SWCNT fluorescence responses of the (8,6), (9,4), and (10,2) chirality peaks were fitted using the Hill isothermal model^[83]^ (**Figure S4**):

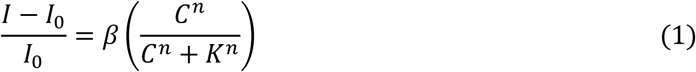

where *I*_0_and *I* are the fluorescence intensities before and after analyte addition, respectively, 𝐶 is the analyte concentration in the solution, 𝐾 is the analyte concentration producing 50% of the maximal fluorescence response, 𝑛 is the Hill coefficient, and 𝛽 is a proportionality factor (**Table S1**). The limit of detection (LOD) was calculated according to the calibration fit for the (9,4) chirality, as the analyte concentration at which the Hill function fit equals three times the standard deviation of the DNA-SWCNT intensity without the analyte, and was found to be 13 nM, 12 nM, and 582 nM for DA with DNA1-SWCNT, TCh with DNA2-SWCNT, and 5-HT with DNA3-SWCNT, respectively.

### 2.2 Spatiotemporal Fluorescence Imaging

To capture spatiotemporal changes in SWCNT fluorescence in response to analytes, fluorescence imaging was conducted with immobilized SWCNTs within PLL-treated microfluidic channels (**Scheme 1A**). The immobilized DNA-SWCNTs were exposed to the respective analyte, followed by a wash, in a controlled, cyclic manner using two automated and synchronized syringe pumps to alternate analyte and PBS flows through the channel **(Scheme 1B**), with four cycles in total. Each cycle started with an 8-minute PBS wash at a flow rate of 900 µL·min^-1^ either to equilibrate the channel in the first cycle or to remove residual analyte in subsequent cycles (**Scheme 1C**). As the microfluidic setup was assumed to maintain laminar flow in the channel, analyte dispersion perpendicularly to the flow direction relied solely on diffusion. Accordingly, for DNA1-SWCNT and DNA2-SWCNT, we introduced the DA and TCh analytes, respectively, for 3 min at the same rate to reach the syringe bulk concentration throughout the channel. Then, the pumps were paused for 3 min to verify that no additional fluorescence changes occurred due to further diffusion, ascertaining complete equilibration. For DNA3-SWCNTs, which displayed distinct kinetics as described below, we extended the 5-HT analyte-flow and no-flow segments to 5 and 10 min, respectively.

**Scheme 1.**
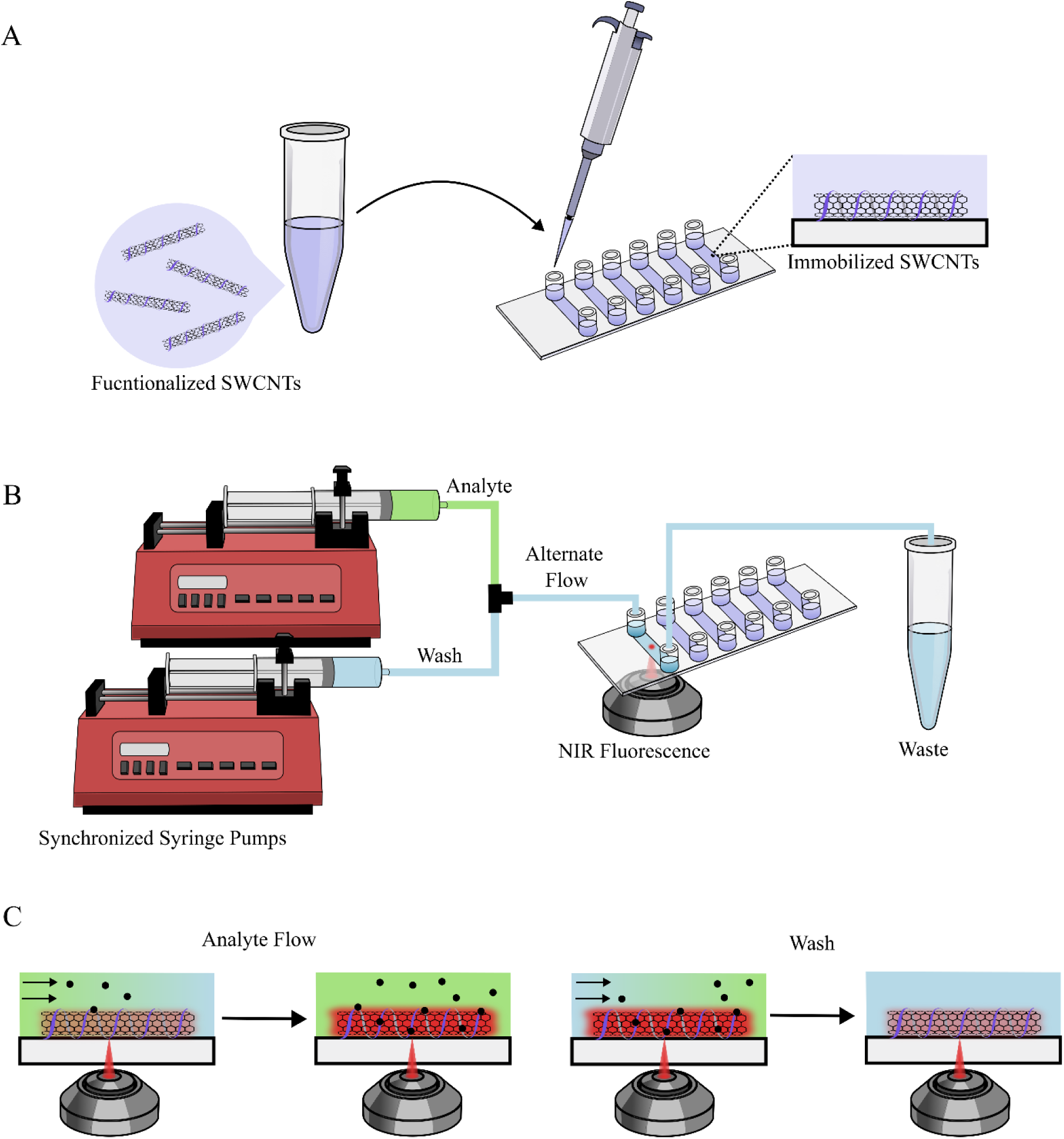
Experimental setup. (A) Functionalized SWCNTs immobilized in microfluidic channels. (B) The immobilized SWCNTs were exposed to the analyte and washed in a cyclical manner using two synchronized and automated syringe pumps with alternating flow, and their NIR fluorescence was recorded using a fluorescence microscope coupled to a NIR camera. (C) During analyte flow, the analyte was uniformly distributed within the channel, interacting with the immobilized SWCNTs and modulating their fluorescence. During the wash step, the analyte molecules were removed from the channel to recover the original SWCNT fluorescence.

Figure 1 presents representative fields of view (FOVs) of NIR fluorescence images before and after analyte addition for each DNA-SWCNT across the four cycles, and the corresponding mean FOV fluorescence intensity time traces. **Figure S5** overlays the mean FOV fluorescence time traces of the four cycles in the flow experiment for each analyte concentration. The three DNA-functionalized SWCNTs exhibited different behaviors throughout the cycles, most pronounced at higher analyte concentrations. DNA1-SWCNT and DNA2-SWCNT both exhibited an immediate increase in fluorescence upon analyte introduction into the channel, followed by a gradual decrease during PBS wash as the analyte was cleared from the channel (Figure 1A-D). Baseline intensities prior to analyte addition, *I*_0_, were largely stable across cycles, with an upward drift at higher analyte concentrations (≥20 µM DA and ≥100 µM TCh) (**Figure S5**A-B, **Table S2**), consistent with incomplete washout and residual analyte on the SWCNTs. The post-analyte fluorescence, *I*, declined in later cycles at higher analyte concentrations for both sensors, but the underlying causes differed. For DNA1-SWCNT, at ≥20 µM DA, the post-analyte intensity decreased both across cycles and, within a given cycle, even while DA continued to flow and during the subsequent no-flow period. This behavior implicates time- and concentration-dependent DA polymerization.^[47,84]^ In contrast, DNA2-SWCNTs maintained stable post-analyte fluorescence within cycles, and a reduction between cycles at ≥10 µM TCh, which can likely be explained by diminished sensitivity after repeated exposures, since TCh shows negligible degradation over hours in neutral buffer.^[85]^

**Figure 1.**
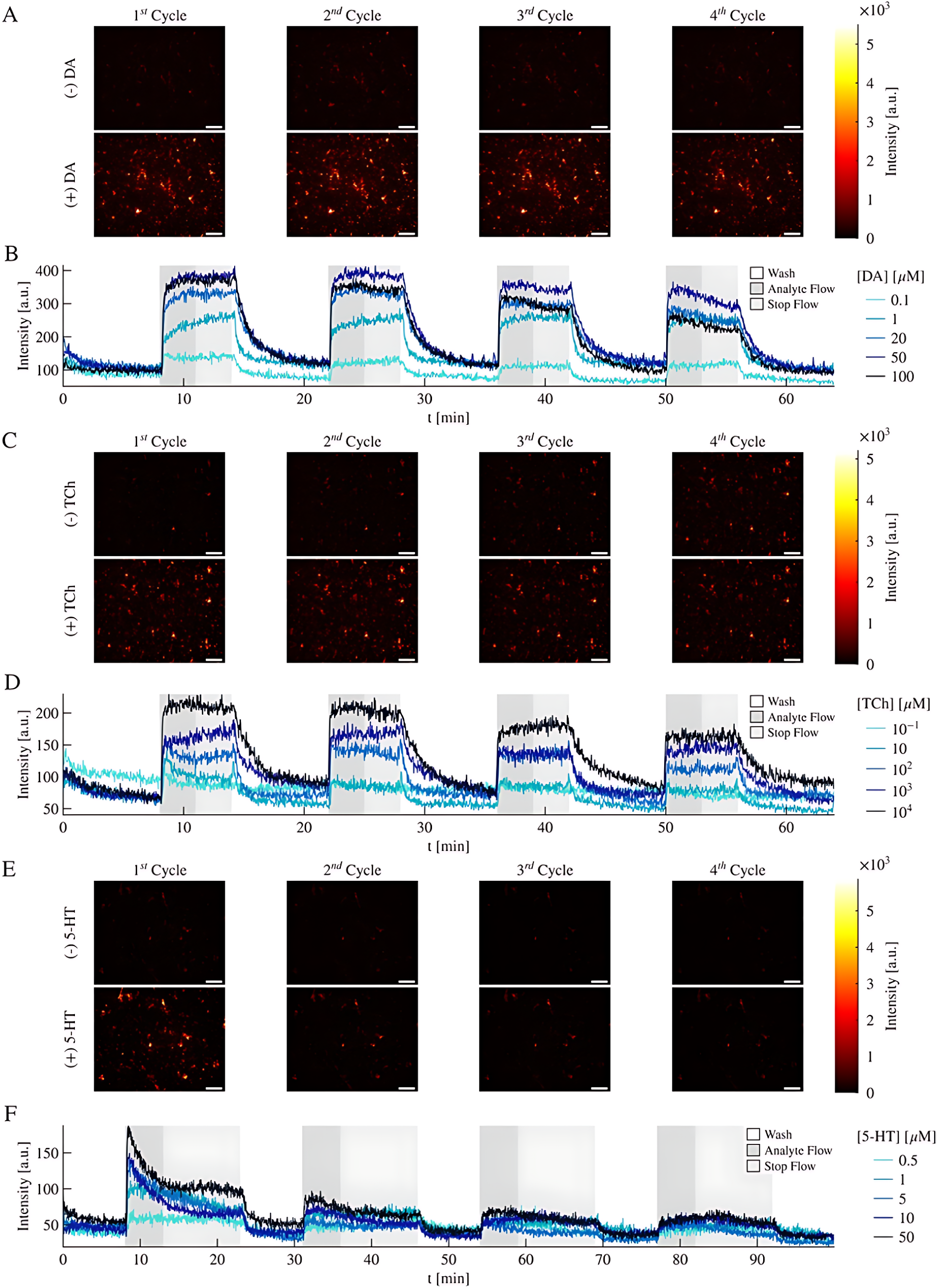
(A) NIR Fluorescence images of the FOV of DNA1-SWCNTs before (top row) and after (bottom row) the addition of 100 µM DA during flow experiments for the four cycles. Scale bar = 10 µm. (B) Mean fluorescence value of the FOV throughout flow experiments of DNA1-SWCNT with DA as an analyte. (C) NIR Fluorescence images of the FOV of DNA2-SWCNTs before (top row) and after (bottom row) the addition of 10 nM TCh during flow experiments for the four cycles. Scale bar = 10 µm. (D) Mean fluorescence value of the FOV throughout flow experiments of DNA2-SWCNT with TCh as an analyte. (E) NIR Fluorescence images of the FOV of DNA3-SWCNTs before (top row) and after (bottom row) the addition of 50 µM 5-HT during flow experiments for the four cycles. Scale bar = 10 µm. (F) Mean fluorescence value of the FOV throughout flow experiments of DNA3-SWCNT with 5-HT as an analyte. White background represents PBS flow, dark grey represents analyte flow, and light grey represents the stop of the pump’s flow for panels B, D, and E.

DNA3-SWCNT exhibited a distinct kinetic profile in response to ≥ 1 µM 5-HT analyte, compared to the two other DNA-SWCNTs. In the first cycle, fluorescence increased upon analyte introduction, then declined within the same cycle before the analyte flow was stopped. To rule out diffusion-limited equilibration as the cause of the fluorescence drop, we extended the analyte-flow period to 5 min. Moreover, to visualize the effect of the subsequent PBS wash under steady-state conditions, we extended the stop-flow interval to 10 min so the FOV stabilized before the next wash, ensuring any additional decrease would be attributed to washing alone (Figure 1F, **Figure S5**C). While 5-HT can undergo polymerization similarly to DA, its polymerization process is significantly slower, spanning days,^[86,87]^ and therefore unlikely to cause the diminished fluorescence. To corroborate that the transient decay is intrinsic to the DNA3-SWCNT–5-HT interaction, we performed a continuous NIR spectroscopy assays of DNA3-SWCNTs in solution phase with the addition of either 50 µM of 5-HT or PBS as a control (**Figure S6**). Both traces showed a brief spike at the moment of addition, an expected artifact from pipette insertion, after which the PBS control remained stable at baseline. In contrast, the 5-HT sample displayed short-lived fluctuations attributable to diffusion and mixing within the well, followed by an increase and subsequent decrease in fluorescence in the absence of any flow. This rise-then-fall profile mirrors the behavior observed for the immobilized DNA3-SWCNTs (Figure 1F), confirming that the post-analyte decay originates from specific SWCNT-analyte interaction rather than from the microfluidic experimental protocol. Comparing with subsequent cycles, DNA3-SWCNTs show a modest decrease in the pre-analyte baseline, *I*_0_, and a pronounced reduction in post-analyte fluorescence, *I*, after the first cycle (Figure 1E-F, Figure 5C **Table S2**).

The normalized SWCNT fluorescence response of the entire FOV was calculated using two normalization schemes based on (*I* − *I*_0_)/*I*_0_. In both, *I* was defined as the mean fluorescence during the first minute after analyte introduction in a given cycle. For the baseline, we either used (i) a cycle-specific baseline *I*_0_, defined as the mean fluorescence over the last minute of PBS wash immediately preceding that cycle’s analyte exposure, or (ii) a fixed baseline 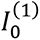, which is the corresponding PBS wash fluorescence intensity value from the first cycle, applied to all cycles. The cycle-specific *I*_0_approach treats each cycle as an independent experiment and facilitates comparison of behavior across cycles when baselines drift. The fixed 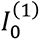 assesses reversibility of *I* relative to the original pre-analyte state, mimicking practical imaging in biological samples, where the true baseline after subsequent exposures may be ambiguous.

To evaluate how immobilized SWCNTs perform under repeated challenges, we quantified the normalized FOV response across cycles and concentrations using the two baselines, namely, the cycle-specific *I*_0_ (Figure 2A-C) and fixed first-cycle 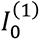 (**Figure S7**A-C). For DNA1-SWCNTs and DA with the cycle-specific baseline *I*_0_, responses are relatively stable for ≤20 µM DA but decline for 20-100 µM in later cycles (Figure 2A). Using the fixed baseline 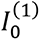 changes this trend, where for lower DA concentrations, the responses appear less consistent across cycles, whereas the responses at higher concentrations change less (**Figure S7**A). Taken together, these results indicate that near and beyond saturation, *I* is comparatively stable (except for a drop in the last cycle with 100 µM DA, attributed to DA polymerization) while *I*_0_ drifts upward with cycles, consistent with residual DA. At low DA concentrations, on the other hand, the dominant effect is variations in *I*, which are less likely to occur when the binding sites on SWCNT are saturated with analyte.

**Figure 2.**
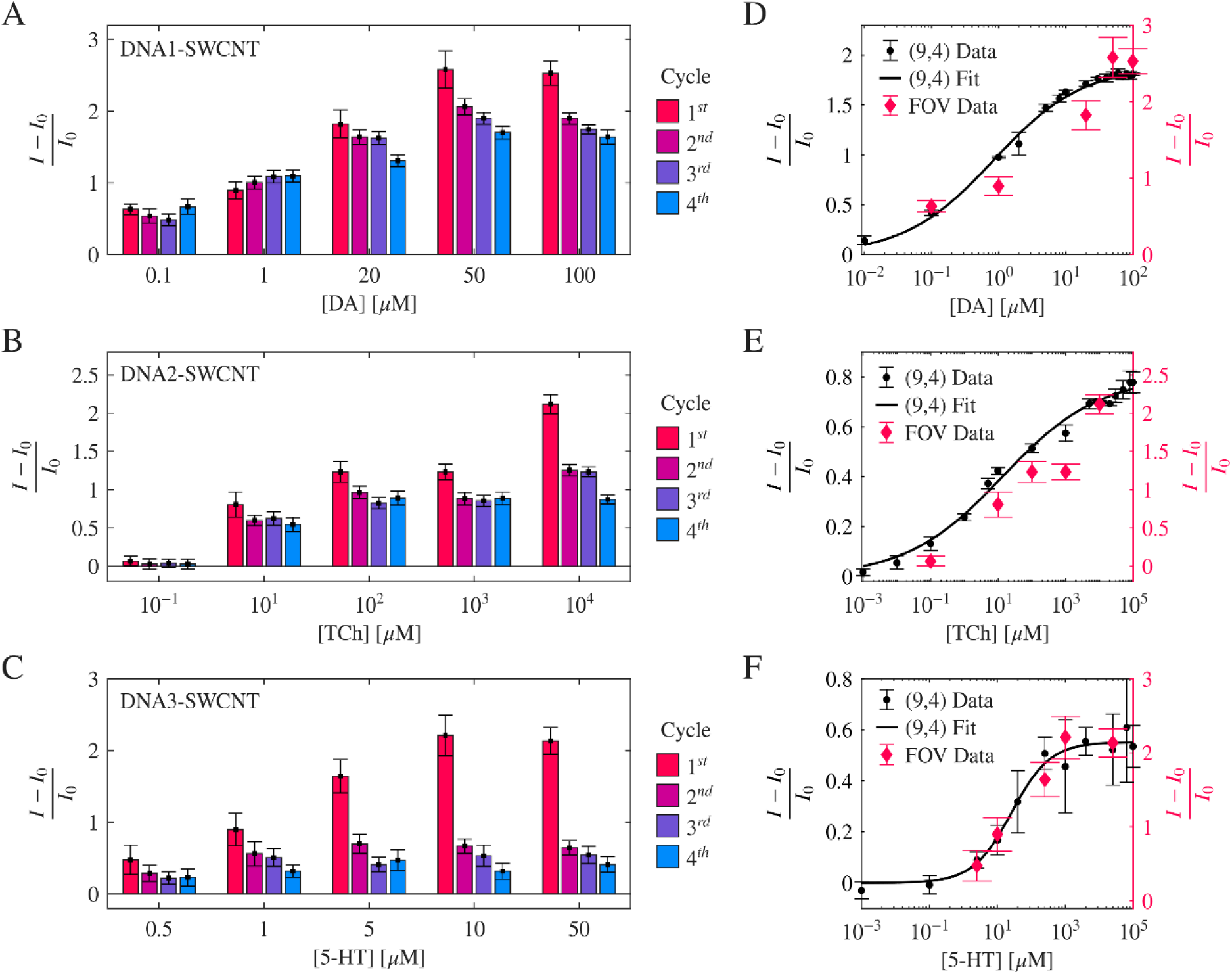
Normalized response of DNA-SWCNT FOV for the different analyte concentrations and cycles, where the normalized response was calculated with both *I* and *I*_0_ values from the current cycle. (A) DNA1-SWCNT with DA as an analyte. (B) DNA2-SWCNT with TCh as an analyte. (C) DNA3-SWCNT with 5-HT as an analyte. (D) Normalized response of DNA-SWCNT FOV from the first cycle compared to the (9,4) chirality calibration curve from Figure S4 for DNA1-SWCNT with DA as an analyte. (E) DNA2-SWCNT with TCh as an analyte. (F) DNA3-SWCNT with 5-HT as an analyte.

The responses of DNA2-SWCNTs to TCh showed similar trends to those of DNA1-SWCNT using the two methods of response calculation, where higher analyte concentrations show better consistency when using a fixed baseline 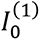, (Figure 2B, Figure S7B). For the 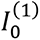 analysis at ≥100 µM, the response in the second cycle is consistent with the first but decreases in the later cycles (Figure S7B), supporting a progressive loss of sensor sensitivity after repeated exposures rather than baseline drift or analyte degradation, as TCh is stable within the experimental timeframe. For lower TCh concentrations, cycle-specific *I*_0_ results in a more consistent response for all cycles, as with DNA1-SWCNT, suggesting a more substantial change in *I* values for lower analyte concentrations.

For DNA3-SWCNTs and 5-HT, the responses using the same-cycle *I*_0_ show a significant loss of response after the first exposure for all 5-HT concentrations (Figure 2C), consistent with the intrinsic rise-then-fall kinetics of this sensor-analyte pair at the first cycle. Holding the baseline fixed using 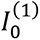 yields a similar trend with an even more pronounced reduction in fluorescence response for all concentrations (**Figure S7**C), indicating that the dominant effect is a decrease in *I* after the first cycle, not baseline drift.

To benchmark immobilized sensors against ensemble behavior in solution phase, we compared the first-cycle imaging response at increasing analyte concentrations with bulk calibration obtained from fluorescence spectroscopy of the dominant (9,4) chirality (Figure 2D-E). Across all three DNA-SWCNT - analyte pairs, the concentration-response curves from imaging closely follow the bulk trends, indicating that immobilization and microfluidic handling preserve the fundamental sensing dynamics on initial exposure. The absolute amplitudes of, however, differ by a scale factor. In bulk, the response is extracted from a specific chirality peak (9,4), whereas imaging integrates emission over the entire field of view from a heterogeneous mixture of chiralities and uses different collection optics and detectors. These methodological differences shift the response scale without altering its shape. Notably, the agreement in concentration dependence trends demonstrates that DNA-SWCNTs retain their functionality after immobilization, validating first-cycle imaging as a faithful proxy for bulk measurements.

### 2.3 Individual regions of interest (ROIs)

Monitoring immobilized SWCNTs enables the detection of localized biochemical events with high spatiotemporal resolution, which is a central motivation for single-sensor imaging in tissues, cell cultures, and *in vivo* applications. Our platform is designed to rigorously characterize single-sensor behavior, including response magnitude, recovery, and stability, thereby laying the groundwork for extracting quantitative information from spatiotemporal imaging. To identify individual SWCNTs, we utilized the iterative thresholding plugin in Fiji, which scans multiple thresholds above a user-defined minimum and assigns each object the threshold at which its segmented area is most stable. Using this approach, SWCNTs-containing regions of interest (ROIs) were detected for each flow experiment (Figure 3A). For the DNA1-SWCNT, DNA2-SWCNT, and DNA3-SWCNT, between 423-623, 279-563, and 140-366 ROIs, representing 4.23-7%, 2.37-5.5%, and 1.22-5% of the entire FOV were detected, respectively. After segmentation, we extracted the mean fluorescence intensity for every ROI over the entire time trace (Figure 3A i, ii, and iii). ROIs whose intensity at any time fell below three times the pre-analyte FOV standard deviation (noise floor) were classified as washed away and excluded from further analysis. This workflow provides a robust, standardized basis for comparing single-sensor responses across analytes, concentrations, and cycles, ultimately aiming to improve quantitative interpretation in high-resolution imaging studies.

**Figure 3.**
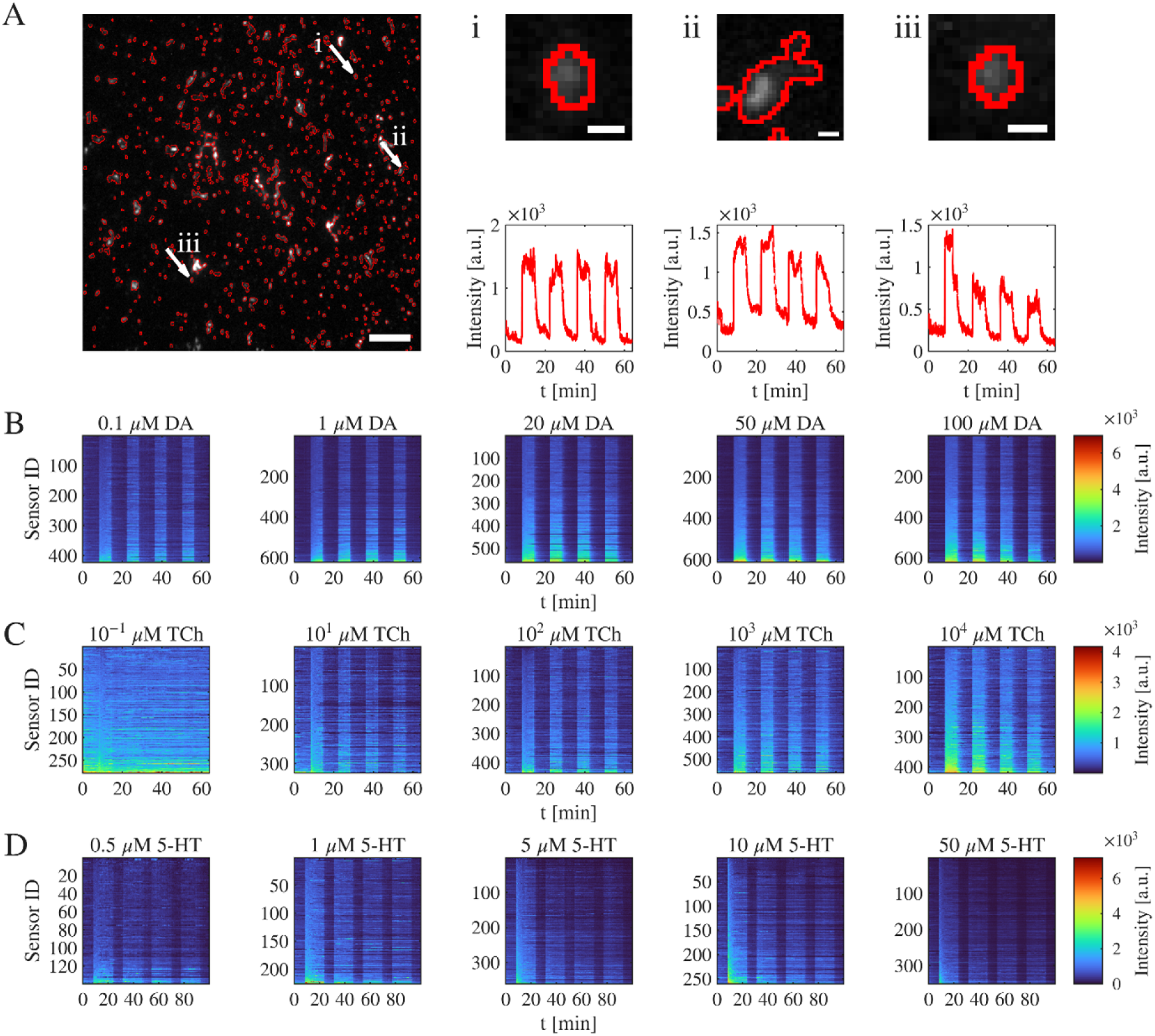
(A) Segmented ROIs of individual SWCNTs in flow experiments of (GT)_15_-SWCNT with 100 µM DA (Scale bar = 10 µm), with (i, ii, iii) three individual ROIs (scale bar = 500 nm) and their mean fluorescence intensity over time. (B) Mean fluorescence intensity traces of individual ROIs across all cycles for all analyte concentrations for DNA1-SWCNTs and DA as an analyte, (C) DNA2-SWCNTs and TCh as an analyte, and (D) DNA3-SWCNTs and 5-HT as an analyte.

Examination of individual ROIs highlights the intrinsic heterogeneity of the SWCNT population (Figure 3A). Nanotube ROIs can vary in length, size, and morphology, where additional diversity arises from mixed chiralities that are indistinguishable in the NIR images, and even within a single chirality, differences in the adsorbed corona conformation further contribute to this diversity.^[88]^ Inspecting the mean fluorescence traces of representative ROIs across cycles (Figure 3Ai-iii) reveals distinct behaviors. Some ROIs exhibit a consistent post-analyte fluorescence, *I*, throughout cycles (Figure 3Ai), while others show a substantial increase in pre-analyte fluorescence baseline, *I*_0_, after the first cycle (Figure 3Aii), indicative of incomplete analyte washout. Additionally, some ROIs exhibited a decrease in *I* values after the first cycle exposure (Figure 3Aiii), which could be attributed to reduced sensitivity to the analyte upon repeated challenges. These observations reveal the variability in both the magnitude and reversibility of the fluorescence response across individual ROIs, which we further explore in subsequent sections.

Despite the variability in ROI fluorescence, when observing the fluorescence time trace of individual ROIs (Figure 3B-D), the fluorescence profile of the ROI populations closely tracks the FOV fluorescence behavior. All ROIs show an increase in intensity upon analyte introduction and a decrease during the PBS wash. Even in cases with minimal ensemble response, e.g., DNA2-SWCNT at 0.1 µM TCh, the ROI population still shows a discernible rise upon TCh addition, especially in the first cycle (Figure 3C). These observations confirm that iterative segmentation indeed identifies SWCNTs, as evidenced by their stimulus responsiveness, and that the ROI ensemble is sufficiently large to capture the overall behavior observed at the FOV level.

With a representative population of SWCNT ROIs in hand, we next quantified single-ROI fluorescence responses to each analyte. For every ROI, we computed the normalized response using both a cycle-specific baseline, *I*_0_ (Figure 4), and a fixed first-cycle baseline 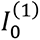 (**Figure S8**), as defined for the FOV analysis. Across all DNA-SWCNTs, analyte concentrations, and cycles, the ROI responses span a wide range, with wider distributions at higher analyte concentrations. Notably, the ROI area alone cannot explain this variability, as the Pearson correlations between the area and first-cycle responses range from -0.097 to 0.332 across conditions, indicating weak correlation (**Figure S9**). Instead, intrinsic heterogeneity is the more plausible driver, specifically, differences in SWCNT chirality, which were also manifested in the chirality-dependent response amplitudes in bulk calibrations, with increasing variability for higher analyte concentrations (**Figure S4**), and in corona conformations that can vary even among nanotubes of the same chirality. The consequence of this heterogeneity is substantial overlap between the response distributions for neighboring concentrations, which limits quantitative inference at true single-sensor resolution. To disentangle the respective contributions of chirality and corona structure, and to possibly tighten the response distributions, studies may employ single-chirality SWCNT samples and more controlled corona architectures.

**Figure 4.**
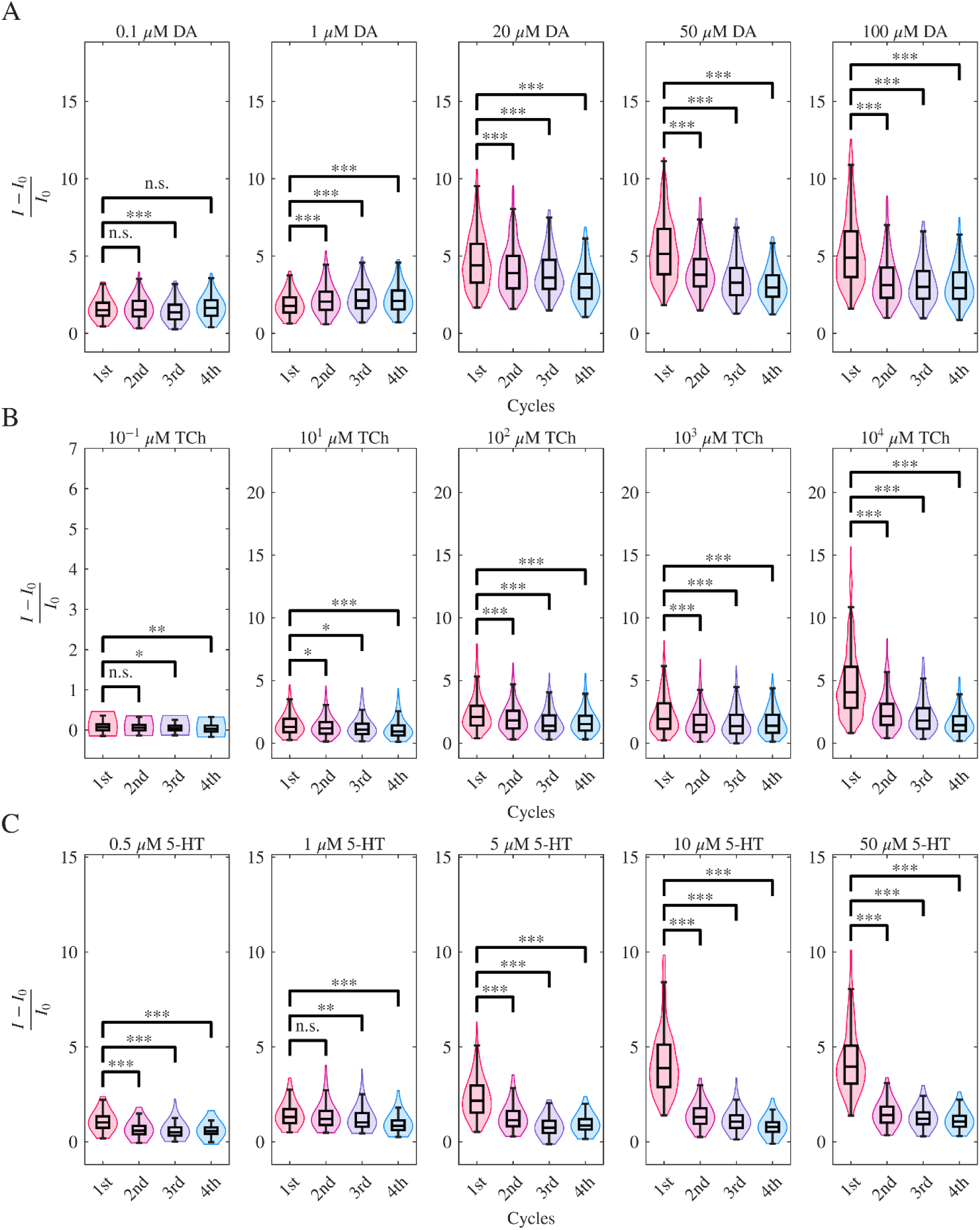
Distribution of the normalized fluorescence response (*I* − *I*_0_)/*I*_0_ to the corresponding analyte across cycles in the segmented ROI population for each DNA-SWCNT and analyte concentration, where *I* and *I*_0_ values both taken from the same cycle. (A) DNA-SWCNT with DA as an analyte. (B) DNA2-SWCNT with TCh as an analyte. (C) DNA3-SWCNT with 5-HT as an analyte. Statistical significance was analyzed using one-way ANOVA tests, n.s.p>0.05, *p<0.05, **p < 0.01, ***p < 0.001.

**Figure 5.**
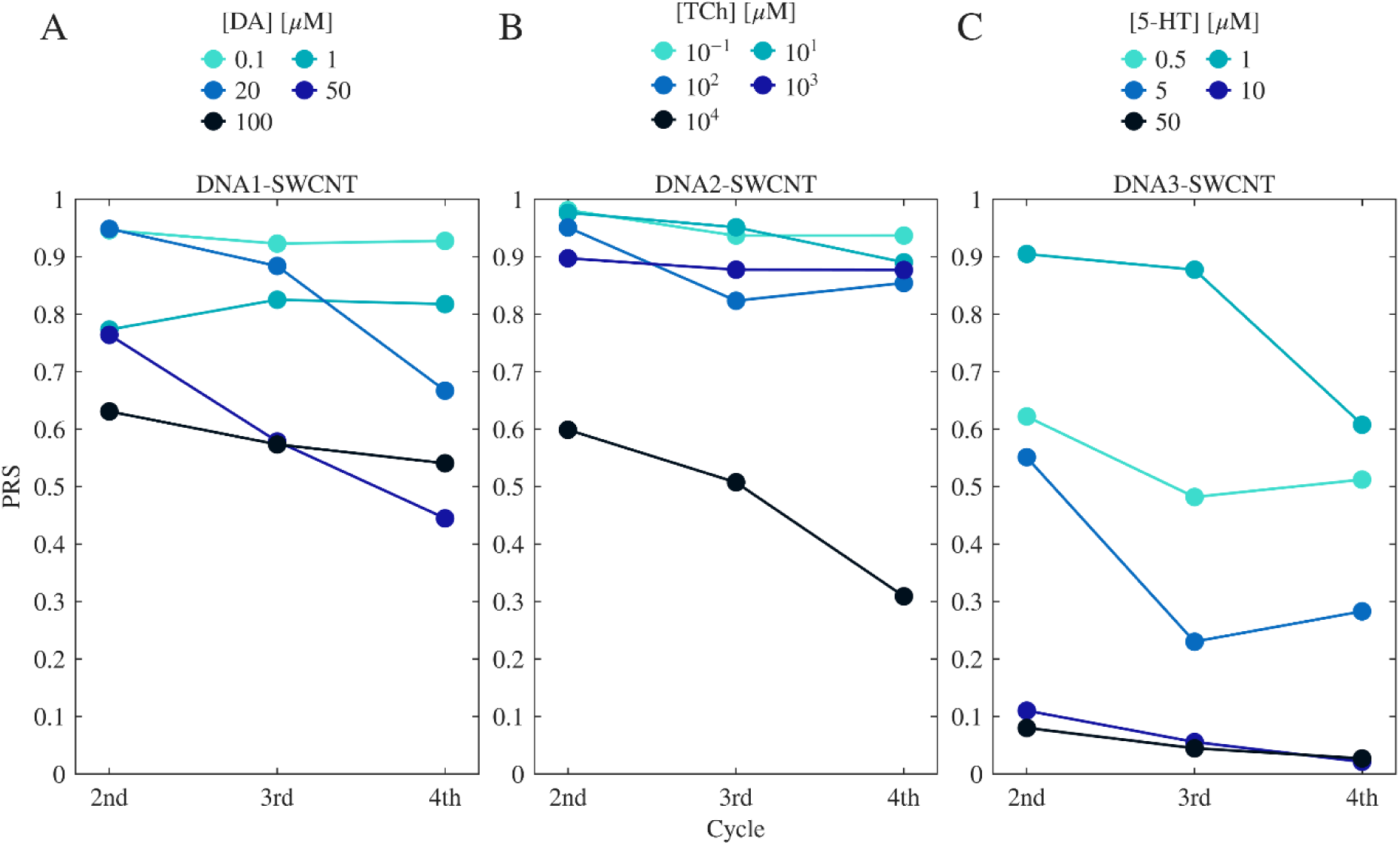
Reversibility Score of normalized fluorescence response, (*I* − *I*_0_)/*I*_0_, of the DNA-SWCNTs to varying analyte concentration across exposure cycles, using cycle-specific baseline *I*_0_. From left to right: DNA1-SWCNTs with DA as an analyte, DNA2-SWCNT with TCh as an analyte, DNA3-SWCNT with 5-HT as an analyte.

The ROI-level response distributions computed with both baselines, cycle-specific *I*_0_ (Figure 4) and fixed 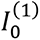 (**Figure S8**), track the same trends observed at the FOV level (Figure 2, **Figure S7**), further confirming that the segmented ROIs constitute a representative SWCNT population of the entire FOV.

We then used one-way ANOVA to assess reversibility by testing for statistically significant differences in ROI responses between the first cycle and later cycles.

DNA1-SWCNTs showed a significant difference between first and later cycles for most of the conditions (Figure 4A and **Figure S8**A). For the responses calculated with 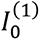, some of the distributions had a lower significance level, or none, compared to the *I*_0_ case (the 2^nd^ and 3^rd^ cycles of 20 µM, all cycles for 50 µM, and the 2^nd^ cycle for 100 µM) indicating that at high DA concentrations, ≥20 µM, *I* remains comparatively stable across cycles, and the apparent variability is dominated by baseline drift, as observed for the entire FOV, with DA polymerization affecting later cycles for ≥20 µM. At low DA levels of 0.1 µM, changes in response in the second cycle are more pronounced than calculated with 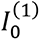, suggesting *I* value changes after analyte exposure, which is further explored in subsequent sections.

For *I*_0_-based responses of DNA2-SWCNT, statistical differences were weaker at ≤10 µM TCh and stronger at ≥100 µM (Figure 4B). Using 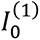, increased significance at low TCh levels but somewhat decreased the significance for the high concentrations, primarily in the second cycle (**Figure S8**B). This suggests that higher TCh concentrations yield more consistent *I* across cycles, whereas lower concentrations show larger cycle-to-cycle changes in *I*.

In the case of DNA3–SWCNT with 5-HT, responses were highly significant (p<0.001) for all concentrations and cycles in both normalization schemes, except the second cycle at 1 µM when using *I*_0_ (Figure 4C, **Figure S8**C). This pervasive significance indicates poor recovery of both *I* and *I*_0_ after the first 5-HT exposure, consistent with the intrinsic rise-then-fall kinetics and subsequent loss of responsiveness described earlier.

### 2.4. SWCNT Reversibility

When using immobilized SWCNTs to monitor analyte presence with high spatial resolution, the same SWCNT can be used repeatedly, undergoing multiple exposures to analyte. The reversibility of SWCNT is therefore fundamental for reliable long-term sensing. To further investigate the reversibility of individual SWCNT and SWCNT populations, we compared *I*_0_ values of individual ROIs in later and their first-cycle baseline 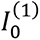 (**Figures S10-S12**). For each ROI, we also estimated the baseline noise as the standard deviation of the pre-analyte trace, and points whose *I*_0_ shift exceeded this noise were flagged in red, indicating changes not attributable to system fluctuations. Data points that fell within this noise level were plotted in green.

For DNA1-SWCNTs and DNA2-SWCNTs, at lower analyte concentrations (≤1 µM DA or ≤1000 µM TCh, respectively), ROIs showed both increases and decreases in *I*_0_ relative to 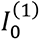 (**Figures S10-S11**).

At higher concentrations, the *I*_0_ values overwhelmingly increase from the original 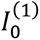 across the ROI populations. Prior work shows that ssDNA coronas (including (GT)₁₅) can reconfigure upon analyte interaction, and SWCNT chirality also influences ssDNA conformation, which can alter the local dielectric environment and charge transfer near the SWCNT, thereby affecting the SWCNT fluorescence.^[47,88–92]^ *I*_0_ shifts at low concentrations could stem from corona remodeling with analyte exposure, as the original corona conformation may not be fully restored following analyte removal. ROI-specific upward or downward *I*_0_shifts support this assumption, as the different corona configurations of SWCNT chiralities could result in different remodeling of the corona after analyte wash, causing differing effects on SWCNT fluorescence. At higher concentrations, however, the near-universal increase in *I*_0_suggests a uniform effect on SWCNT corona after analyte wash, which is consistent with incomplete analyte removal (residual DA or TCh).

DNA3-SWCNT ROIs, in contrast, exhibit a consistent decrease in *I*_0_ values after the first cycle of 5-HT exposure, for all concentrations and subsequent cycles (**Figure S12**). Together with the unique rise-and-fall behavior of DNA3-SWCNT during the first analyte exposure (Figure S5C), this suggests that the first analyte challenge induces a substantial change in SWCNT functionalization and corona, which in turn decreases the SWCNT fluorescence during the first cycle and limits subsequent recovery afterwards.

We next compared changes in post-analyte intensity *I* to changes in baseline *I*_0_ values for individual ROIs between later and first cycles. To better understand the relationship between the changes in fluorescence values across cycles, a linear regression was performed, and the R^2^ value and the slope of the regression were calculated (**Figures S13-S15**). For DNA1-SWCNT and DNA2-SWCNT, the R^2^ and slope values present an overall decreasing trend with increasing analyte concentrations. This suggests that at lower analyte concentrations, the changes in fluorescence values between cycles better adhere to a linear relationship (**Figures S13-S14**). This behavior, together with the observations of the changes in *I*_0_ values (**Figures S10-11**), suggests that for lower analyte concentrations, the changes in *I* values between cycles are linearly dependent on *I*_0_ changes. If baseline shifts arise from analyte-induced corona reconfiguration, this linearity suggests that changes in *I* values might also be caused by changes in the corona, implying the response mechanism of the SWCNTs remains intact while the system “resets” to a new baseline. This explains the better apparent reversibility when responses are normalized to the cycle-specific *I*_0_ rather than the fixed 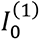 at low concentrations (Figure 4A-B, **Figure S8**A-B), and it implies that reliable quantification under repeated low-dose exposures benefits from updating *I*_0_after each challenge. For higher analyte concentrations, the linear relationship between *I* and *I*_0_ changes is less pronounced, with smaller 𝑅^2^ and slope values, as *I* is relatively consistent while *I*_0_ increases. As previously discussed, the increase in *I*_0_ values is likely due to incomplete washing of analyte molecules from the SWCNTs. The relatively consistent *I* values may reflect the lack of “reset” of baseline corona conformation when analyte remains bound, or saturation of binding sites at high analyte concentrations, which reduces variability in *I* values. For both DNA1-SWCNT and DNA2-SWCNT, there is a uniform decrease in *I* values of the ROIs across cycles for higher analyte concentrations (≥20 µM DA, ≥10 µM TCh). While the effect in DNA1-SWCNT is likely due to DA polymerization, as discussed previously, for DNA2-SWCNT, the effect is not attributable to analyte degradation, thus could be explained by a decreased sensitivity after multiple challenges.

For DNA3-SWCNT, the 𝑅^2^ and slope values of the linear regression also exhibit a decreasing trend with increasing 5-HT concentrations (**Figure S15**). However, as described before, the majority of ROIs show a substantial decrease in both *I* and *I*_0_values, especially in higher 5-HT concentrations. This pattern indicates that analyte-induced corona changes are sufficiently strong to irreversibly decrease baseline fluorescence and reduce the sensitivity of SWCNTs, compromising reversibility in subsequent exposures.

### 2.5 Population Reversibility Score

To summarize recovery behavior in a single, comparable metric, we defined a Population Reversibility Score (PRS) applicable to any model of sensor-analyte pair. Since the immobilized sensor population is heterogeneous, comprising mixtures of chiralities and variations in corona conformation (even within a given chirality), length, and surface-adsorption morphology, the PRS reflects ensemble recovery. We quantify reversibility directly from the distribution of single-sensor responses in each experiment. For every cycle and concentration, the ROI responses are converted into a probability density using kernel density estimation (KDE), a non-parametric method that estimates the probability density function of a random variable based on a finite set of data samples.^[93]^ We then measure the statistical distance to a reference distribution, namely, the first-cycle distribution at the same concentration, using the Kullback-Leibler (KL) Divergence (relative entropy):^[94]^

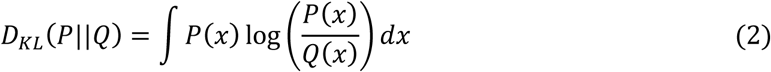

where *P* and *Q* denote the cycle-specific and reference response distributions, respectively. The relative entropy is a non-negative measure that quantifies how one probability distribution diverges from another, and it equals zero only when the two distributions are identical. Thus, it increases as the cycle’s distribution departs from its first-cycle counterpart, capturing changes arising from any source of variability within the heterogeneous population. To map this unbounded distance to an intuitive, unitless score in the range of [0,1], we define the *PRS* to be:

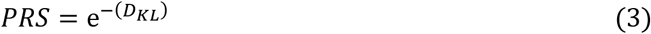

By construction, *PRS* = 1 indicates perfect recovery (the cycle’s distribution matches the first cycle), whereas *PRS* values approaching 0 indicate poor reversibility. This cycle- and concentration-dependent *PRS* compresses the full population statistics into a single number, enabling standardized comparisons across constructs, doses, and experimental conditions.

We computed the PRS using both normalization schemes, cycle-specific *I*_0_ (Figure 5) and fixed 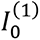 (**Figure S16**), to assess how reversibility depends on analyte concentration, number of exposures, and baseline choice. At lower analyte concentrations, all three DNA-SWCNTs achieve higher *PRS* when responses are normalized to the cycle-specific baseline *I*_0_, consistent with previous observations of the stronger linearity in change of *I*_0_ and *I* values between cycles for lower analyte concentrations, possibly due to corona confirmation change after the first exposure. At higher concentrations, the trend flips for DNA1-SWCNTs (DA) and DNA2-SWCNTs (TCh), where PRS is higher with the fixed baseline 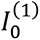, indicating that the post-analyte intensity *I* remains comparatively stable while *I*_0_drifts (e.g., due to residual analyte). DNA3-SWCNT (5-HT) exhibits lower PRS scores for fixed baseline response, likely due to its unique rise and fall response dynamics, which caused a reduced baseline and sensitivity of the SWCNTs after the first 5-HT exposure. Beyond baseline choice, the PRS captures the cumulative impact of repeated challenges. For most concentrations and all SWCNT constructs, PRS decreases with cycle number, revealing a practical limit to the number of exposure-wash cycles the sensors can undergo before reversibility degrades.

The *PRS* highlights functionalization-dependent differences in SWCNT reversibility and, therefore, should be assessed for any sensor intended for long-term imaging to judge whether consecutive measurements are valid and comparable. For sensors with high *PRS* across multiple exposures, sequential readouts from long-term imaging could be considered reliable for comparisons and concentration estimates. However, when *PRS* is low or declines with cycling, quantitative use requires prior multi-cycle calibration (or limiting the number of exposures), as the response cannot be assumed to be reversible. Beyond flagging stability, *PRS* also indicates the preferred normalization strategy (e.g., cycle-specific *I*_0_ vs. fixed 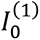) and guides experimental choices on whether to re-baseline between cycles, limit analyte concentration ranges, strengthen wash protocols, or cap the number of exposure-wash rounds.

## 3. Conclusion

In this work, we established a controlled microfluidics imaging-based platform to quantify how optical nanosensors respond and recover under repeated analyte challenges. The reversibility of the SWCNTs is a critical parameter for their reliable long-term use as fluorescence-based analyte sensors. Notably, first-cycle responses of immobilized sensors tracked ensemble (bulk) calibration, confirming that the core concentration-dependent response dynamics are preserved upon immobilization. Nevertheless, by resolving hundreds of individual sensor ROIs and analyzing pre-analyte and post-analyte fluorescence intensity modulations across multiple analyte exposure cycles, we revealed broad variability in response magnitude and signal recovery, and we linked multi-cycle changes to analyte- and dose-dependent mechanisms. For DNA1-SWNCT and DNA2-SWCNT, at lower analyte concentrations, individual ROIs generally retained their fluorescence response dynamics despite shifts in baseline fluorescence, suggesting that initial analyte exposure induced a change in corona conformation without permanently impairing sensing function. In contrast, higher analyte concentrations led to systematic increases in baseline fluorescence values, likely due to an incomplete analyte washout, limiting sensor recovery. DNA3-SWCNT exhibited a distinct rise-then-fall behavior with irreversible decrease in fluorescence, indicating stronger or more disruptive analyte interactions with the corona layer of the SWCNTs.

A key design choice in this first demonstration was to work with mixed-chirality samples, which posed a greater challenge for quantifying reversibility. The immobilized populations include multiple chiralities, in addition to heterogeneity in length, corona conformation, and surface-adsorbed morphology, which can arise stochastically when casting onto substrates or immobilizing in hydrogels/tissues. This heterogeneity increases the distributional spread and concentration overlap at the single-sensor level, thereby stress-testing any metric that aims to summarize recovery. Although using single-chirality materials can narrow variance, even such samples retain inherent heterogeneity in corona states, lengths, and adsorption geometries, and a population-level measure of reversibility therefore remains necessary.

To standardize reporting across constructs, concentrations, and cycles, we introduced the Population Reversibility Score (PRS), derived from the KL divergence between cycle-specific and first-cycle response distributions. PRS maps distributional differences onto a unitless [0,1] scale, with PRS=1 representing perfect recovery, enabling robust and generalizable comparisons of long-term operational stability. This score accurately summarizes the progressive loss of fluorescence response reversibility across repeated cycles, demonstrating its dependence on both analyte concentration and the number of exposures. Furthermore, the analysis highlighted that the choice of baseline calculation is crucial for maximizing apparent reversibility, where a cycle-specific baseline provides a better PRS for low analyte concentrations, reflecting a true “system reset”, while the fixed first-cycle baseline is better suited for high concentrations.

More broadly, our platform and the Population Reversibility Score are agnostic to the specific SWCNT-analyte pairing and can be extended to other corona chemistries, target molecules, and chirality enrichment strategies. Beyond SWCNTs, the same workflow of microfluidic cycling, single-sensor (or single-ROI) distributional analysis, and KL-based reversibility scoring, applies to any sensing modality that relies on a transient signal to report analyte presence over time. As such, this framework offers a general path to standardize and compare long-term recovery in systems designed for continuous, high-resolution monitoring in complex environments.

## 4. Experimental Section

### Materials

HipCO SWCNTs were purchased from NanoIintegris. Single-stranded DNA sequences were purchased from Integrated DNA Technologies. Dopamine hydrochloride was purchased from Sigma-Aldrich, Thiocholine Iodide from BOC Sciences, and Serotonin (hydrochloride) from Cayman Chemical.

### SWCNT Suspension Preparation

HipCO SWCNTs were dispersed with three ssDNA sequences, (GT)_15_ (DNA1), T_30_ (DNA2), and CCC CCC AGC CCT TCA CCA CCA ACT CCC CCC (DNA3), in NaCl (0.1 M) using bath sonication (Elma P-30H, 80 Hz) followed by tip sonication on ice (Qsonica Q125, 3 mm tip, 4 W). For DNA1 and DNA2, SWCNTs (1 mg) with ssDNA (2 mg) were bath-sonicated for 10 min and tip-sonicated twice for 20 min each, whereas for DNA3, SWCNTs (1 mg) with ssDNA (100 µM) were bath-sonicated for 2 min and tip-sonicated for 10 min, according to the protocol in Kelich *et al*.^[48]^. All suspensions were then centrifuged twice at 16,100 rcf (DNA1 and DNA2 for 90 min per spin, and DNA3 for 30 min per spin), each time collecting the top 80% of the supernatant.

### Absorption Spectroscopy

The absorption of the suspensions was recorded between 200 nm and 1400 nm (1 nm step size) using a UV-vis-NIR spectrophotometer (Shimadzu UV-3600 PLUS). The concentration of the suspension was calculated according to the absorption measured at 632nm, using an extinction coefficient of ε_632nm_ = 0.036 L·mg^−1^·cm^−1^.^[47]^

### Excitation-Emission Spectra

SWCNT solutions (0.5 mg·L^−1^) in PBS (pH 7.4) were illuminated using a supercontinuum white-light laser (NKT-photonics, Super-K Extreme) with a bandwidth filter (NKT-photonics, Varia, Δλ = 20 nm). They were scanned between 450 nm and 840 nm with a 2 nm wavelength step size, at an intensity of 20 mW (at 730 nm). Emission spectra were recorded by a NIR inverted fluorescence microscope (Olympus IX73), and spectrally resolved using a spectrograph (Spectra Pro HRS-300, Princeton Instruments) with a slit-width of 500 μm and a grating (150 g·mm^-1^). An InGaAs camera (PylonIR, Teledyne Princeton Instruments) was used to record the fluorescence intensity spectra.

### NIR Fluorescence Spectroscopy

In a 96-well plate, functionalized SWCNT solutions (0.5 mg·L^-1^) in PBS were added to each well, followed by the addition of analytes in different concentrations. The well plate was mounted on the stage of the inverted microscope (Olympus IX73) and illuminated by a supercontinuum white-light laser, as described above, at an excitation wavelength of 730 nm with an intensity of 20 mW.

### Continuous NIR Fluorescence Spectroscopy

A 96-well plate, with 145 µM of DNA3-SWCNT solution (0.5 mg·L^-1^) in PBS per well, was placed on a mounted stage of the inverted microscope (Olympus IX73), and illuminated by a 730 nm continuous-wave laser (MDL-MD-730 1.5 W, Changchun New Industries) with an intensity of 100 mW. The fluorescence emission was recorded continuously with an exposure time of 2 s and spectrally resolved as described above. After a 1-minute baseline, 5 µL of either PBS or serotonin (final concentration of 50 µM) was added.

### SWCNTs Immobilization

For flow experiments, Ibidi µ-slides VI 0.1 were used. The channels were incubated with PLL (Poly-L-Lysine) (5 µL, 0.01%) for 10 min, washed with ddH2O, and dried. Then, functionalized SWCNT solution (0.5 mg·L^-1^, 5 µL) in PBS was added to the channels for 10 min incubation, washed with PBS, and dried. All channels were filled with PBS (90 µL) to maintain SWCNT hydration until imaging.

### Fluorescence Imaging

The prepared slide was placed on the stage of an inverted fluorescence microscope (Olympus IX83) equipped with a UPLFLN 100X objective lens, 1.3 NA. Two syringes, filled with medium (PBS) and analyte solution, were placed in automated syringe pumps (SyringeTwo, NewEra) and connected to the channel through a tubing system. The medium and analyte were flowed through the channel at a flow rate of 900 µL·min^-1^. During the flow experiment, the SWCNT fluorescence was excited by a 730 nm continuous-wave laser (MDL-MD-730 1.5 W, Changchun New Industries) with an intensity of 500 mW. The laser excitation light was directed to the sample by a dichroic mirror (900 nm lp, chroma, T900lpxxrxt), and the NIR emission of the SWCNTs was detected after an additional 900 nm long-pass emission filter (chroma, ET900lp) with an InGaAs camera (Raptor, Ninox 640 VIS-SWIR).

### Image Processing

All images were processed by Fiji (ImageJ) and MATLAB. Individual SWCNTs in the images were identified using the ‘3D iterative thresholding’ plugin in Fiji, with MSER as the threshold criterion.

## Acknowledgements

G.B. acknowledges the support of the Zuckerman STEM Leadership Program, the ERC NanoNonEq 101039127, the Israel Science Foundation (Grant no. 196/22), the Ministry of Science, Technology, and Space, Israel (Grant no. 1001818370 and 0008452), the Marian Gertner Institute for Medical Nanosystems at Tel Aviv University, the Tel Aviv University Center for AI and Data Science (TAD), the Zimin Institute for Engineering Solutions Advancing Better Lives, and the Naomi Prawer Kadar Foundation.

---

**ToC.**
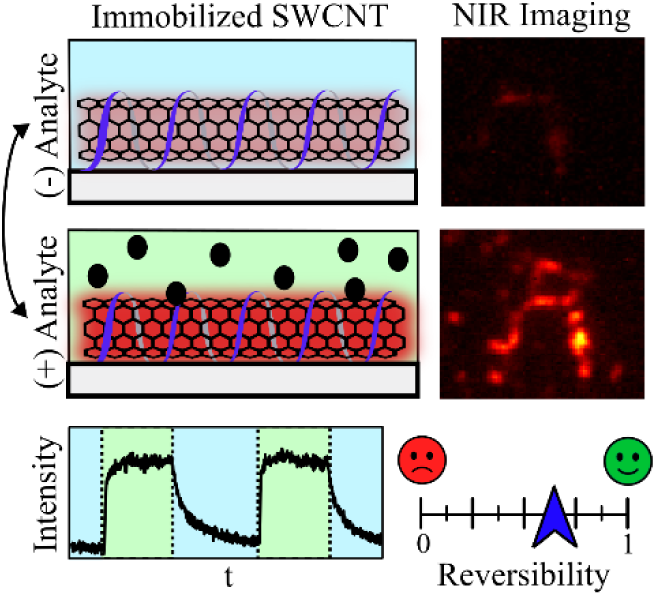
Single-walled carbon nanotubes (SWCNTs) are versatile optical nanosensors that can be imaged at the single-sensor level to map biological processes. To translate such imaging into calibrated measurements, sensor recovery under repeated analyte exposures must be analyzed. This work introduces a multi-cycle assay and a Population Reversibility Score to systematically characterize the reversibility of sensor populations.

