## Supplemental Information for "Quantifying Population Reversibility of Sensor Performance in Multi-Cycle Single-Sensor Recovery Assay"

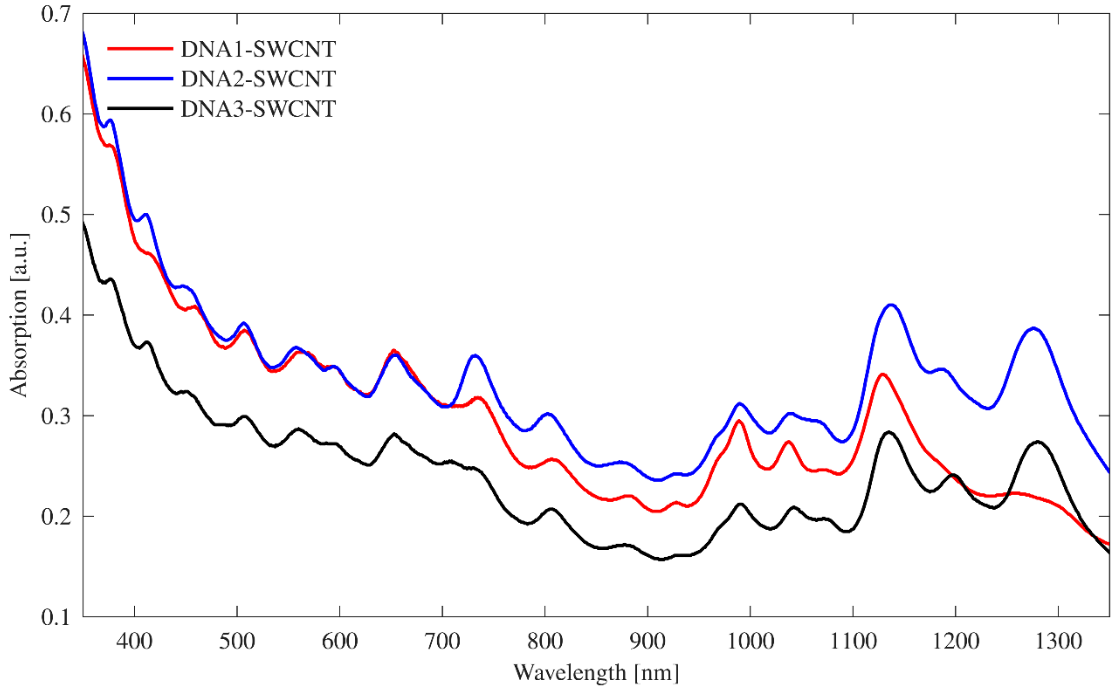


**Figure S1**. Absorption spectrum of ssDNA functionalized SWCNTs.

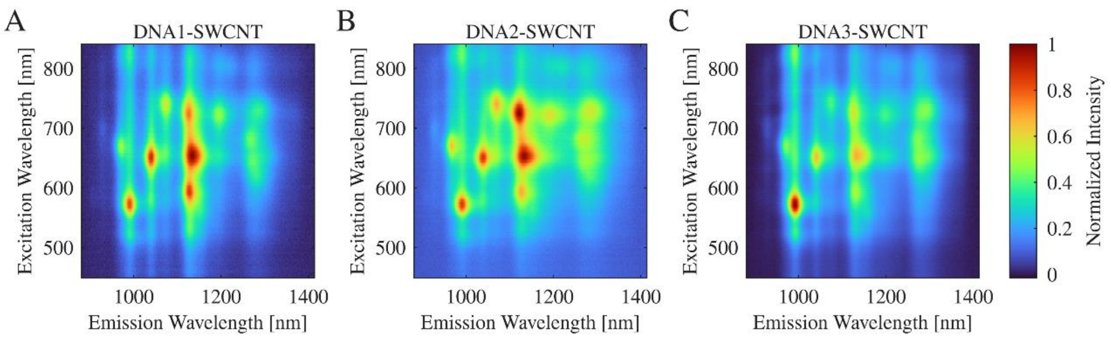


**Figure S2**. Excitation-emission maps normalized to the maximum emission intensity of ssDNA functionalized SWCNT. (A) DNA1-SWCNT. (B) DNA2-SWCNT. (C) DNA3-SWCNT

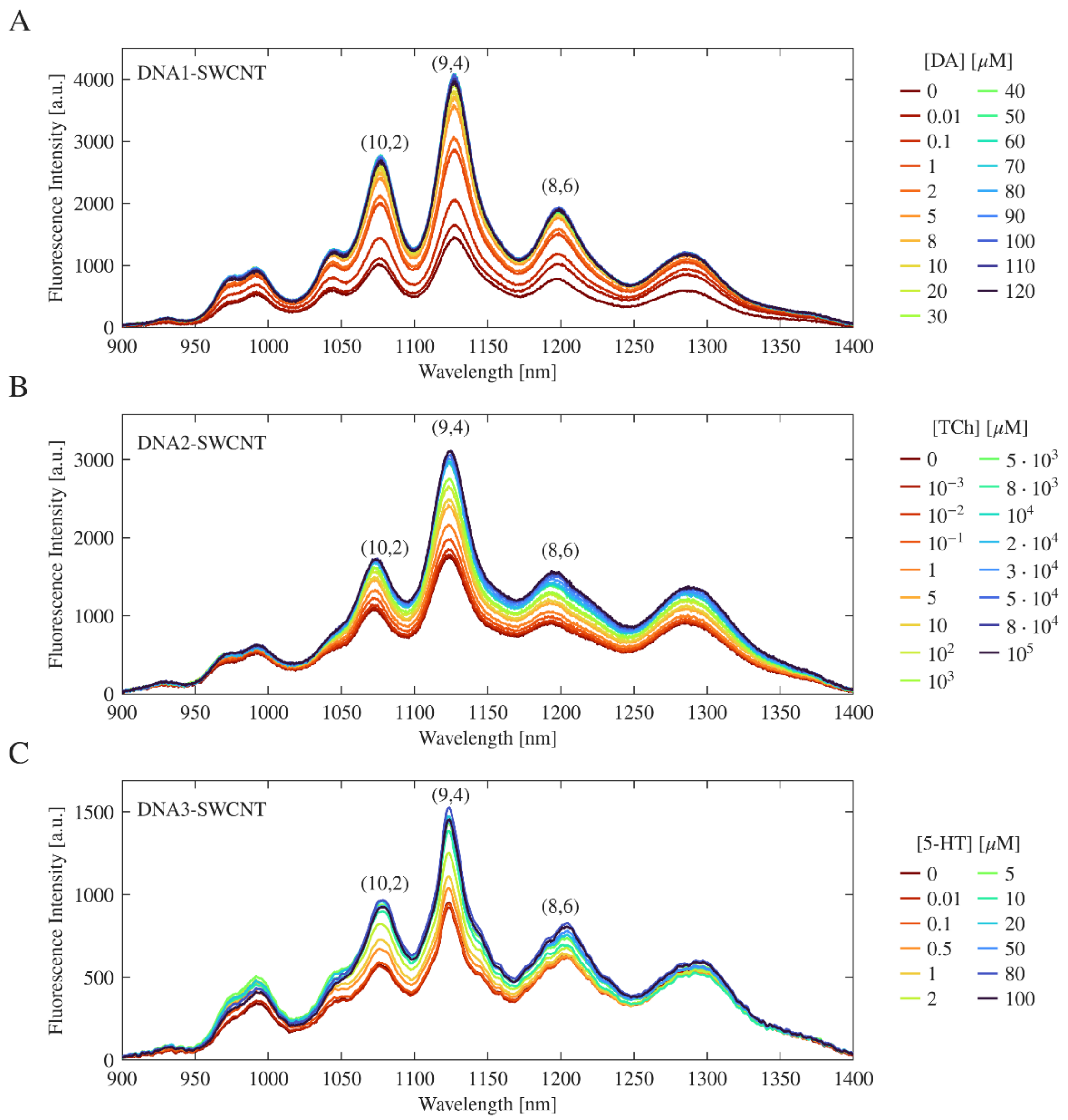


**Figure S3**. Fluorescence spectra of ssDNA functionalized SWCNT upon the introduction of increasing concentrations of analyte. (A) DNA1-SWCNT with the introduction of DA. (B) DNA2-SWCNT with the introduction of TCh. (C) DNA3-SWCNT with the introduction of 5-HT. Chirality peaks of (10,2), (9,4), and (8,6) chiralities are marked.

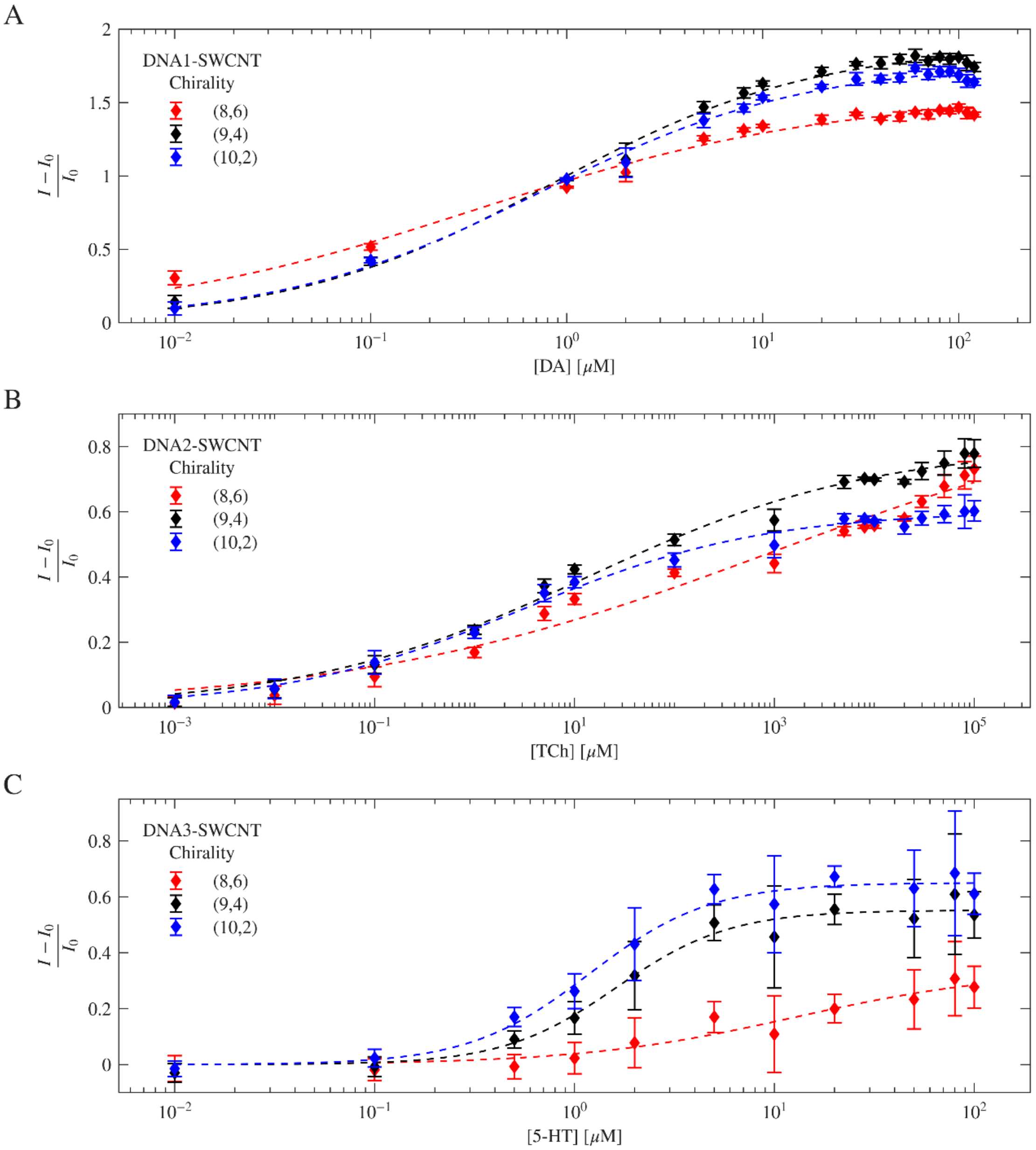


**Figure S4**. Calibration of ssDNA-functionalized SWCNT normalized fluorescence response of different SWCNT chiralities to the corresponding analyte, where the data points represent experimental data ($n = 3$), and the dashed line represents the calibration fit according to the Hill equation. (A) DNA1-SWCNTs response to DA. (B) DNA2-SWCNTs response to TCh. (C) DNA3-SWCNTs response to 5-HT.

**Table S1**. Calibration fit parameters and their 95% confidence intervals, and LOD calculated according to the calibration curve.

| | **Chirality** | **β** | **K [μM]** | **n** | **LOD [μM]** |
|---|---|---|---|---|---|
| **DNA1-SWCNT** | (8,6) | 1.55 (1.48, 1.62) | 0.352 (0.23, 0.47) | 0.481 (0.4, 0.56) | 0.002 |
| | (9,4) | 1.895 (1.83, 1.96) | 0.842 (0.65, 1.03) | 0.652 (0.56, 0.75) | 0.013 |
| | (10,2) | 1.777 (1.72, 1.83) | 0.733 (0.58, 0.73) | 0.641 (0.56, 0.73) | 0.011 |
| **DNA2-SWCNT** | (8,6) | 0.958 (0.54, 1.38) | 1001 (0, 5968) | 0.205 (0.12, 0.29) | 0.022 |
| | (9,4) | 0.801 (0.74, 0.86) | 13.58 (1.6, 25.57) | 0.305 (0.24, 0.37) | 0.012 |
| | (10,2) | 0.603 (0.58, 0.63) | 3.04 (1.58, 4.5) | 0.361 (0.3, 0.42) | 0.048 |
| **DNA3-SWCNT** | (8,6) | 0.339 (0.13, 0.55) | 12.65 (-13.77, 39.07) | 0.807 (0.06, 1.55) | 5.315 |
| | (9,4) | 0.552 (0.51, 0.6) | 1.622 (1.1, 2.14) | 1.532 (0.84, 2.23) | 0.582 |
| | (10,2) | 0.65 (0.61, 0.69) | 1.19 (0.88, 1.5) | 1.437 (0.89, 1.98) | 0.318 |

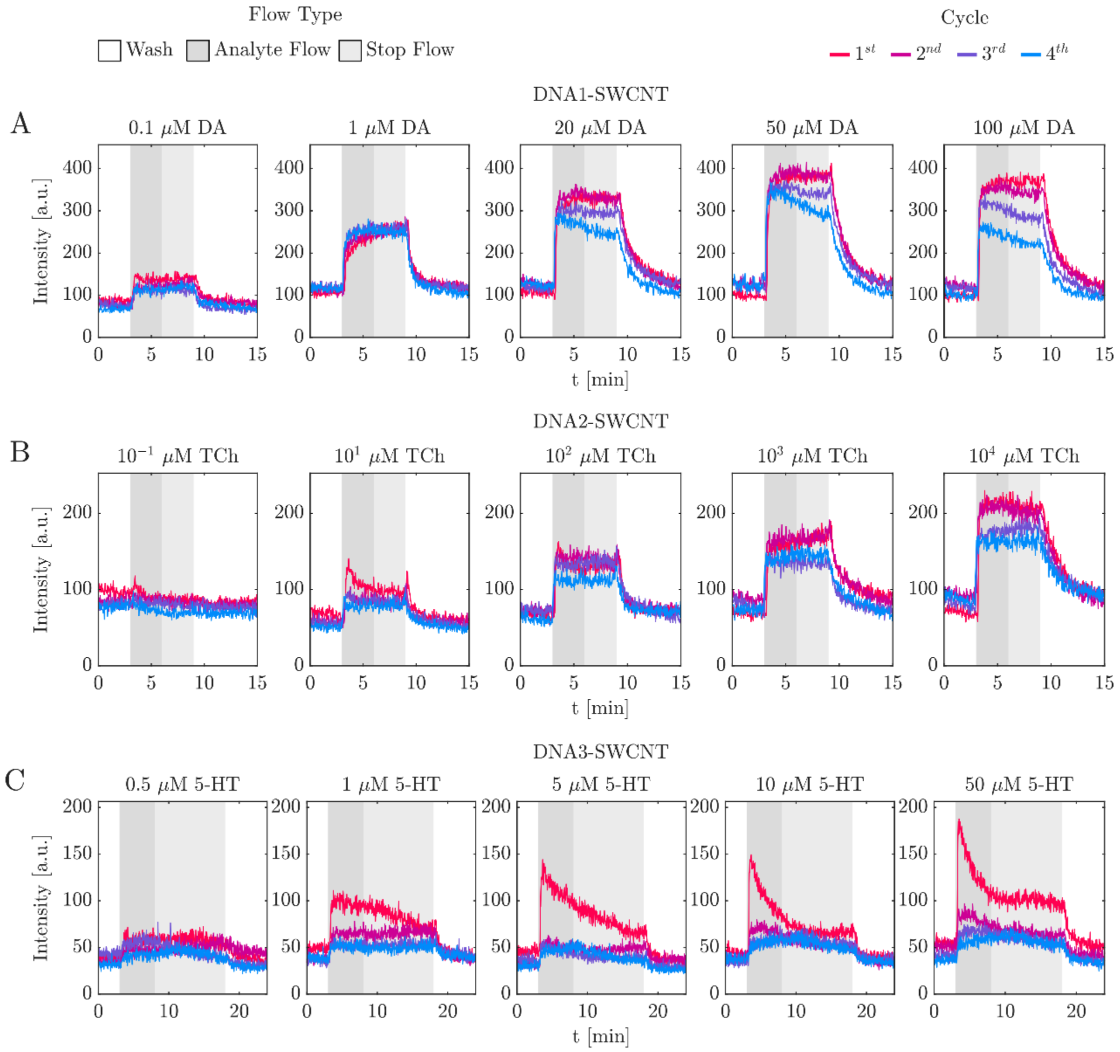


**Figure S5**. Comparison of the mean fluorescence value of the FOV between cycles. White background represents PBS flow, dark grey represents analyte flow, and light grey represents the stop of the pump's flow. (A) DNA1-SWCNT flow experiments with DA as an analyte. (B) DNA2-SWCNT flow experiments with TCh as analyte. (C) DNA3-SWCNT flow experiments with 5-HT as analyte.

**Table S2**. Mean FOV $I_0$ and $I$ values for each SWCNT sensor, analyte concentration, and cycle.

| | [Analyte] [μM] | | 1st Cycle | 2nd Cycle | 3rd Cycle | 4th Cycle |
|---|---|---|---|---|---|---|
| **DNA1-SWCNT** | 0.1 | $I_0$ | 85.7 ± 5.3 | 75.5 ± 5.5 | 72.9 ± 5.9 | 67.8 ± 4.8 |
| | | $I$ | 139.8 ± 6.2 | 116.1 ± 7.4 | 108.3 ± 5.9 | 113.1 ± 7.0 |
| | 1 | $I_0$ | 107.8 ± 5.3 | 112.9 ± 6.6 | 117.2 ± 5.9 | 115.5 ± 6.3 |
| | | $I$ | 204.3 ± 12.9 | 226.2 ± 10.0 | 244.7 ± 10.4 | 242.1 ± 10.0 |
| | 20 | $I_0$ | 105.7 ± 6.7 | 123.2 ± 6.5 | 114.1 ± 7.7 | 120.4 ± 6.3 |
| | | $I$ | 298.1 ± 20.3 | 325.1 ± 12.4 | 299.6 ± 10.0 | 278.1 ± 9.7 |
| | 50 | $I_0$ | 96.8 ± 4.7 | 121.9 ± 8.2 | 121.5 ± 12.6 | 125.7 ± 12.1 |
| | | $I$ | 346.5 ± 25.1 | 373.1 ± 14.0 | 352.4 ± 9.9 | 339.7 ± 11.4 |
| | 100 | $I_0$ | 98.2 ± 5.5 | 119.7 ± 5.3 | 115.3 ± 5.9 | 97.6 ± 7.2 |
| | | $I$ | 346.4 ± 16.3 | 347.1 ± 9.3 | 316.7 ± 7.4 | 257.4 ± 9.6 |
| **DNA2-SWCNT** | 0.1 | $I_0$ | 94.6 ± 5.2 | 84.4 ± 5.4 | 82.6 ± 4.2 | 79.0 ± 4.5 |
| | | $I$ | 100.8 ± 6.0 | 86.7 ± 5.8 | 85.9 ± 4.3 | 81.2 ± 5.1 |
| | 10 | $I_0$ | 65.7 ± 4.1 | 57.1 ± 3.2 | 55.5 ± 3.2 | 50.6 ± 3.1 |
| | | $I$ | 118.7 ± 10.9 | 91.1 ± 3.9 | 90.1 ± 4.9 | 78.2 ± 4.7 |
| | $10^2$ | $I_0$ | 64.8 ± 4.8 | 72.4 ± 4.1 | 72.6 ± 4.1 | 59.5 ± 4.4 |
| | | $I$ | 144.6 ± 8.7 | 142.4 ± 5.8 | 132.5 ± 5.4 | 112.5 ± 5.5 |
| | $10^3$ | $I_0$ | 68.6 ± 3.7 | 86.2 ± 4.4 | 74.7 ± 4.7 | 73.3 ± 5.2 |
| | | $I$ | 153.0 ± 7.2 | 162.3 ± 7.3 | 138.5 ± 5.6 | 138.3 ± 6.1 |
| | $10^4$ | $I_0$ | 67.0 ± 4.2 | 91.1 ± 5.3 | 76.7 ± 4.7 | 86.5 ± 11.0 |
| | | $I$ | 208.8 ± 8.3 | 205.4 ± 6.9 | 171.2 ± 4.9 | 162.0 ± 5.2 |
| **DNA3-SWCNT** | 0.5 | $I_0$ | 38.3 ± 4.1 | 34.3 ± 3.0 | 43.9 ± 3.6 | 31.6 ± 2.4 |
| | | $I$ | 56.6 ± 7.9 | 44.2 ± 3.9 | 53.6 ± 3.8 | 38.9 ± 3.8 |
| | 1 | $I_0$ | 49.4 ± 3.9 | 38.5 ± 2.6 | 34.5 ± 3.0 | 38.1 ± 3.9 |
| | | $I$ | 93.9 ± 11.2 | 60.1 ± 6.5 | 52.0 ± 4.2 | 50.2 ± 3.3 |
| | 5 | $I_0$ | 47.0 ± 4.1 | 30.8 ± 2.8 | 36.2 ± 3.1 | 30.6 ± 2.6 |
| | | $I$ | 124.2 ± 10.9 | 52.4 ± 4.1 | 51.1 ± 3.7 | 45.1 ± 4.4 |
| | 10 | $I_0$ | 41.7 ± 3.5 | 40.1 ± 3.0 | 34.3 ± 2.4 | 38.2 ± 3.2 |
| | | $I$ | 133.9 ± 11.9 | 66.8 ± 4.1 | 52.6 ± 5.0 | 50.3 ± 4.3 |
| | 50 | $I_0$ | 54.2 ± 3.0 | 50.9 ± 3.8 | 41.2 ± 3.6 | 37.4 ± 2.7 |
| | | $I$ | 169.8 ± 10.2 | 83.7 ± 5.2 | 63.8 ± 4.9 | 52.8 ± 4.1 |

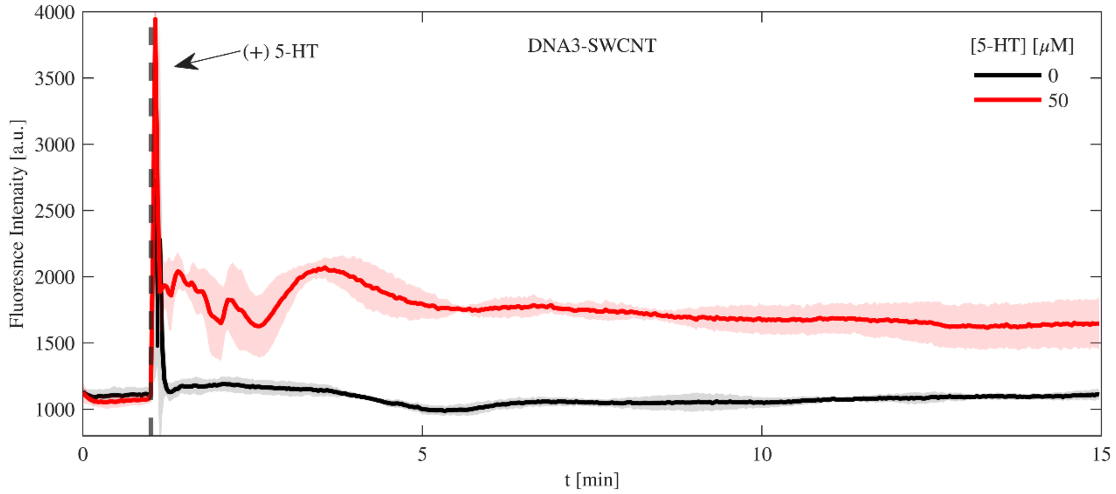


**Figure S6.** Continuous bulk fluorescence response of the (9,4) chirality of DNA3-SWCNT to 5-HT. The red line represents the fluorescence response to 50 µM 5-HT, while the black line represents the control (0 µM 5-HT).

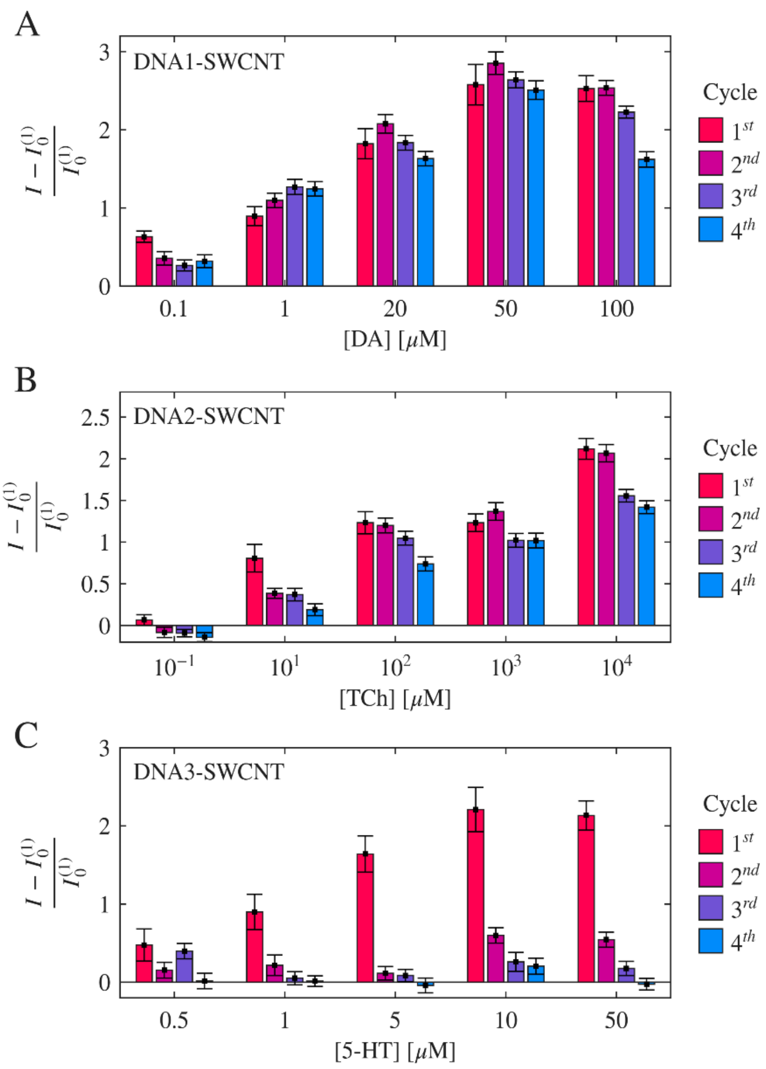


**Figure S7**. Normalized response of DNA-SWCNTs FOV for different analyte concentrations and cycles, where the normalized response is calculated with post-analyte fluorescence values from the current cycle, $I$, and baseline values from the first cycle, $I_0^{(1)}$. (A) DNA1-SWCNT with DA. (B) DNA2-SWCNT with TCh. (C) DNA3-SWCNT with 5-HT.

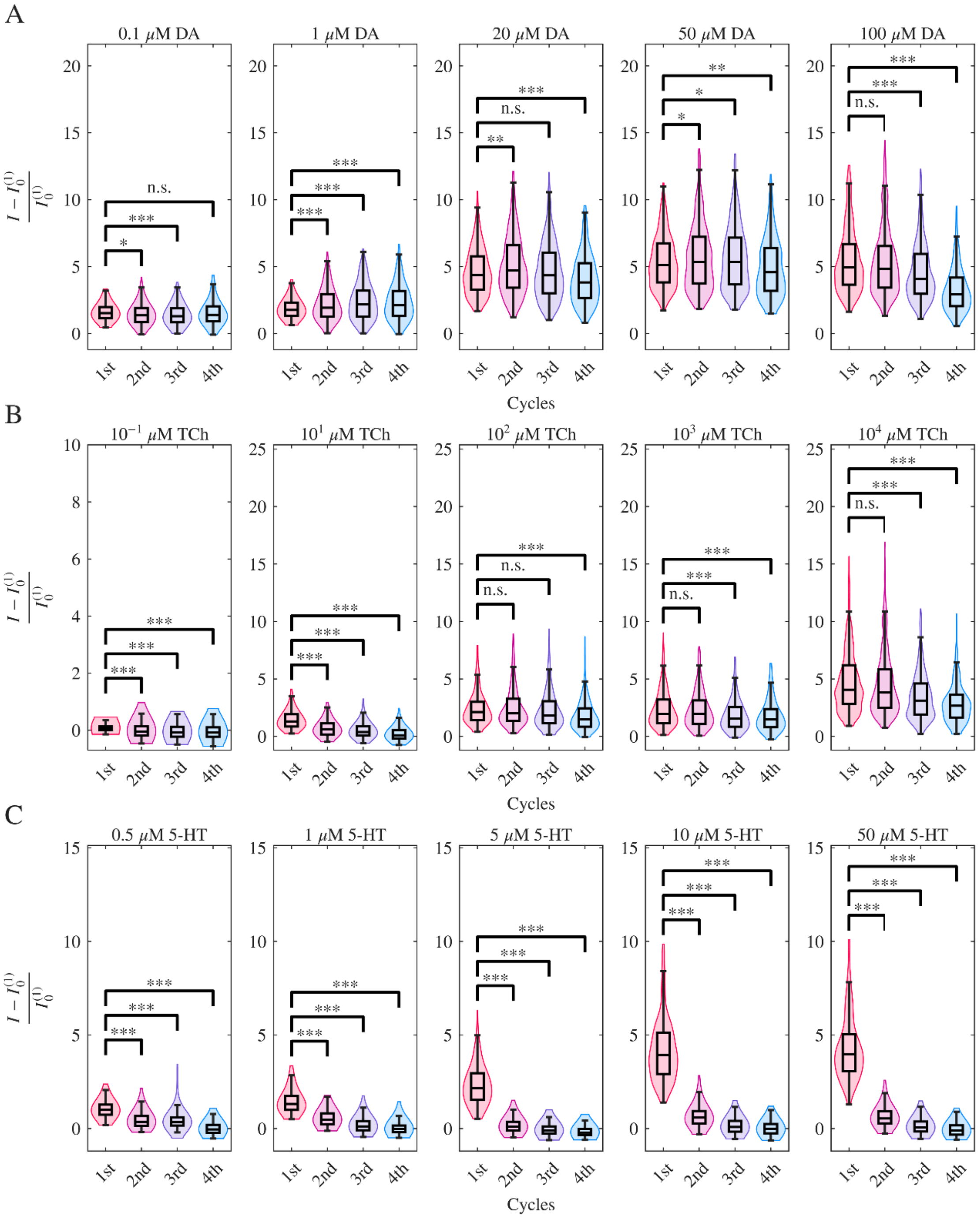


**Figure S8**. Distribution of the normalized fluorescence response $\left(I - I_0^{(1)}\right)/I_0^{(1)}$ to analyte through cycles in the segmented ROI population for each DNA-SWCNT and analyte concentration, where $I$ was taken from the current cycle, and $I_0^{(1)}$ the first cycle. (A) DNA-SWCNT with DA as an analyte. (B) DNA2-SWCNT with TCh as an analyte. (C) DNA3-SWCNT with 5-HT as an analyte. Statistical significance was analyzed using one-way ANOVA tests, n.s. p>0.05, *p<0.05, **p < 0.01, ***p < 0.001.

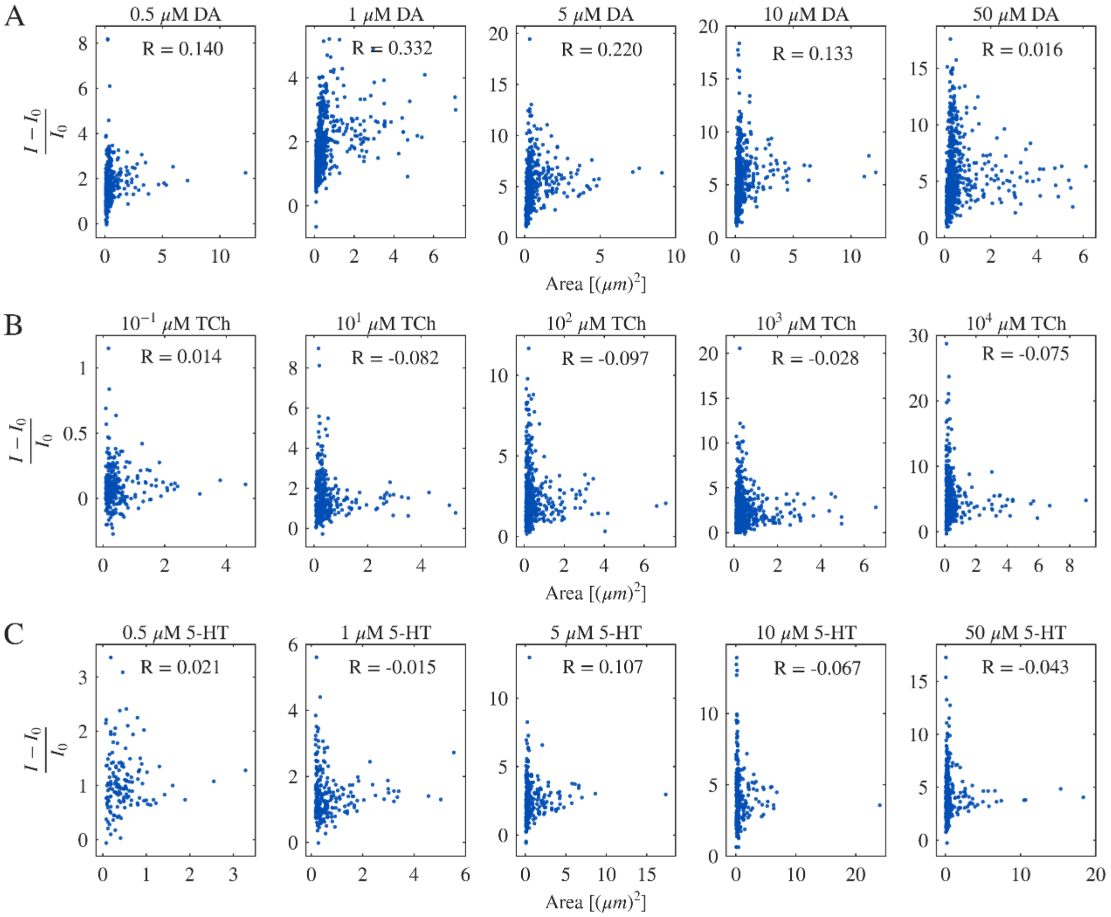


**Figure S9**. Individual ROI normalized fluorescence response $(I - I_0)/I_0$ vs. ROI area in the first cycle of flow experiments with different analyte concentrations. The Pearson coefficient, $R$, between the two variables was calculated to determine linear correlation. (A) DNA1-SWCNT with DA as analyte. (B) DNA2-SWCNT with TCh as analyte. (C) DNA3-SWCNT with 5-HT as analyte. Each dot represents a different ROI.

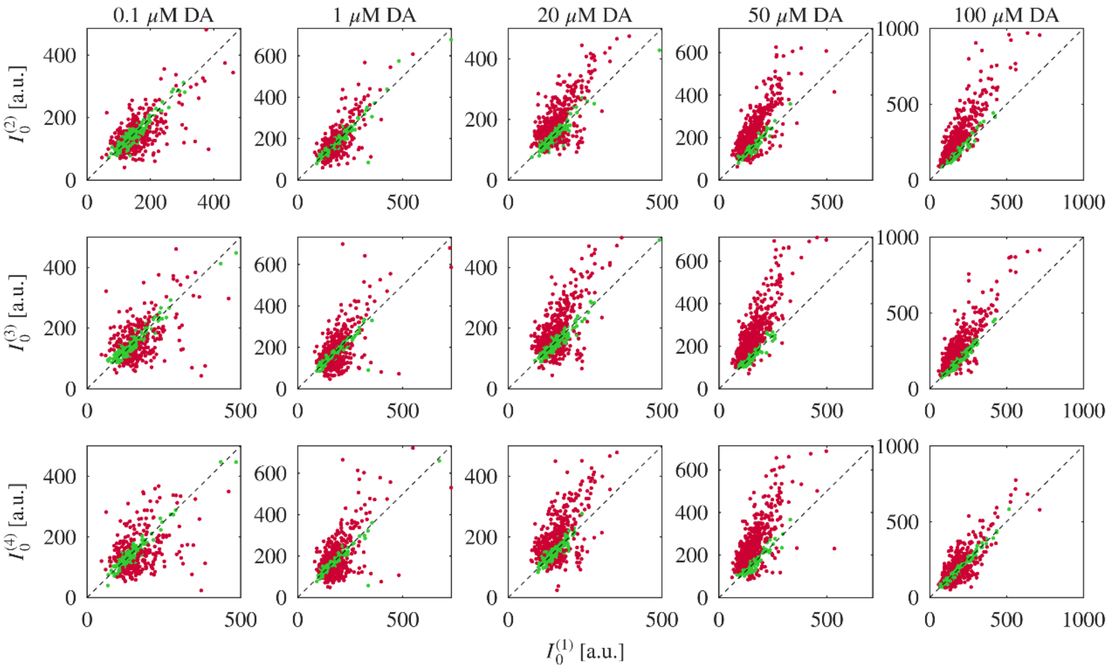


**Figure S10.** Comparison of $I_0$ across cycles for DNA1-SWCNT at varying DA concentrations. Each point is an individual ROI, plotting later-cycle $I_0$ (y-axis) versus first-cycle $I_0^{(1)}$ (x-axis). The dashed line denotes the $x = y$ line to aid visualization of increases (above the line) or decreases (below the line) in $I_0$ relative to the first cycle. Green points indicate ROIs whose $I_0$ shift is within ±1 STD of that ROI's initial baseline cycle $I_0^{(1)}$ (noise range), and red points exceed ±1 STD, indicating a baseline change beyond noise.

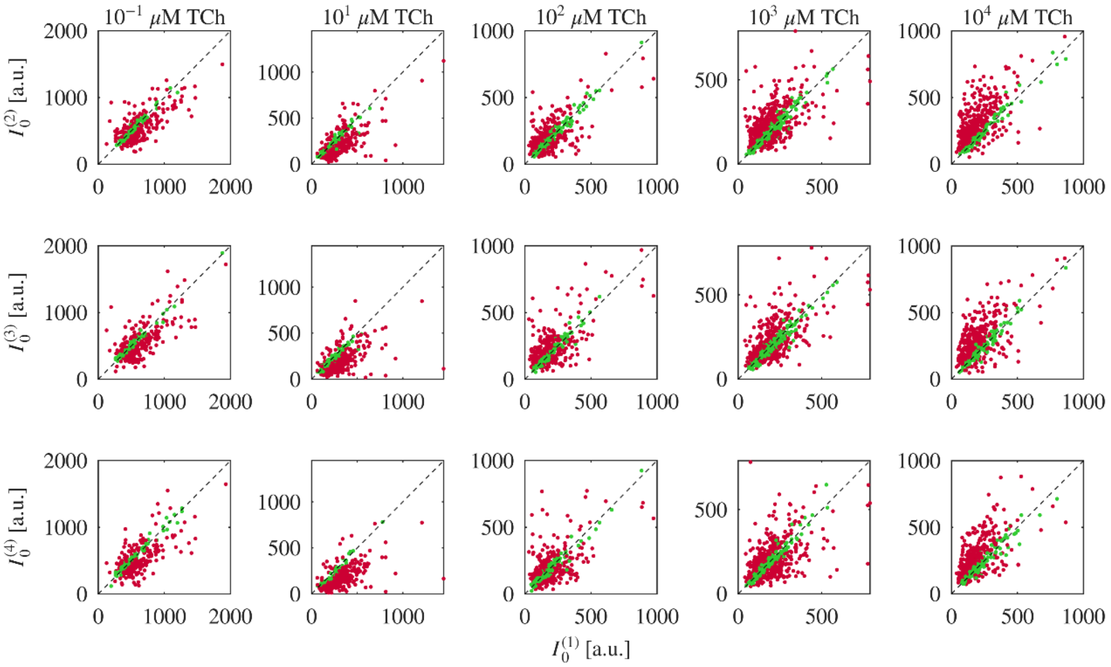


**Figure S11.** Comparison of $I_0$ across cycles for DNA2-SWCNT at varying TCh concentrations. Each point represents an individual ROI, plotting later-cycle $I_0$ (y-axis) versus first-cycle $I_0^{(1)}$ (x-axis). The dashed line denotes the $x = y$ line to aid visualization of increases (above the line) or decreases (below the line) in $I_0$ relative to the first cycle. Green points indicate ROIs whose $I_0$ shift is within ±1 STD of that ROI's initial baseline cycle $I_0^{(1)}$(noise range), and red points exceed ±1 STD, indicating a baseline change beyond noise.

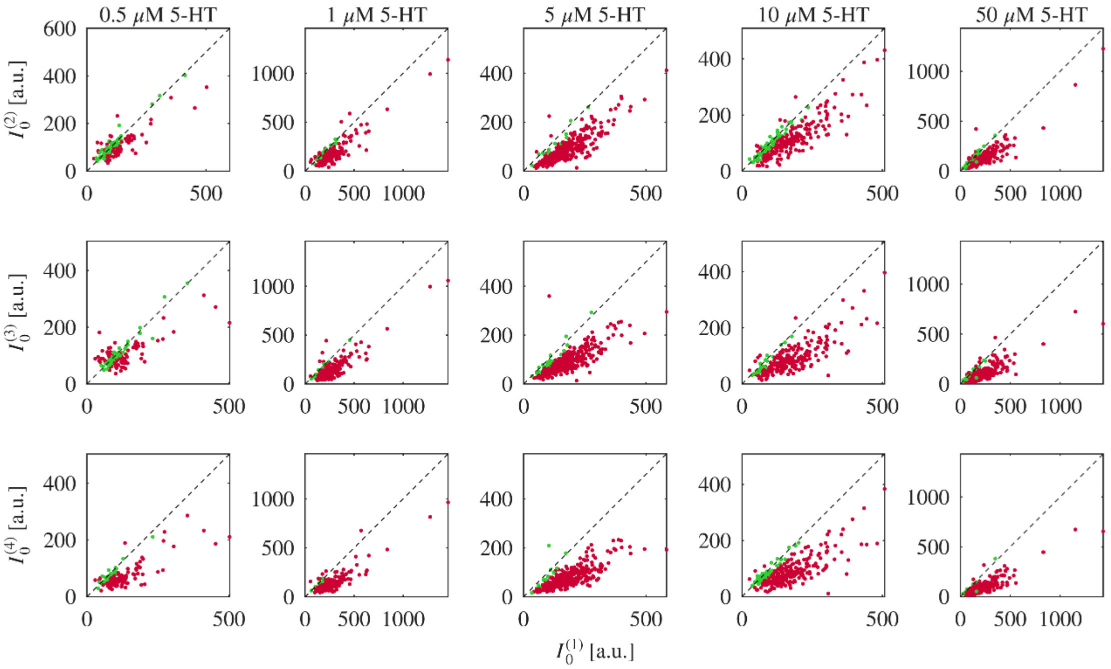


**Figure S12.** Comparison of $I_0$ across cycles for DNA3-SWCNT at varying 5-HT concentrations. Each point is an individual ROI, plotting later-cycle $I_0$ (y-axis) versus first-cycle $I_0^{(1)}$ (x-axis). The dashed line denotes the $x = y$ line to aid visualization of increases (above the line) or decreases (below the line) in $I_0$ relative to the first cycle. Green points indicate ROIs whose $I_0$ shift is within ±1 STD of that ROI's initial baseline cycle $I_0^{(1)}$ (noise range), and red points exceed ±1 STD, indicating a baseline change beyond noise.

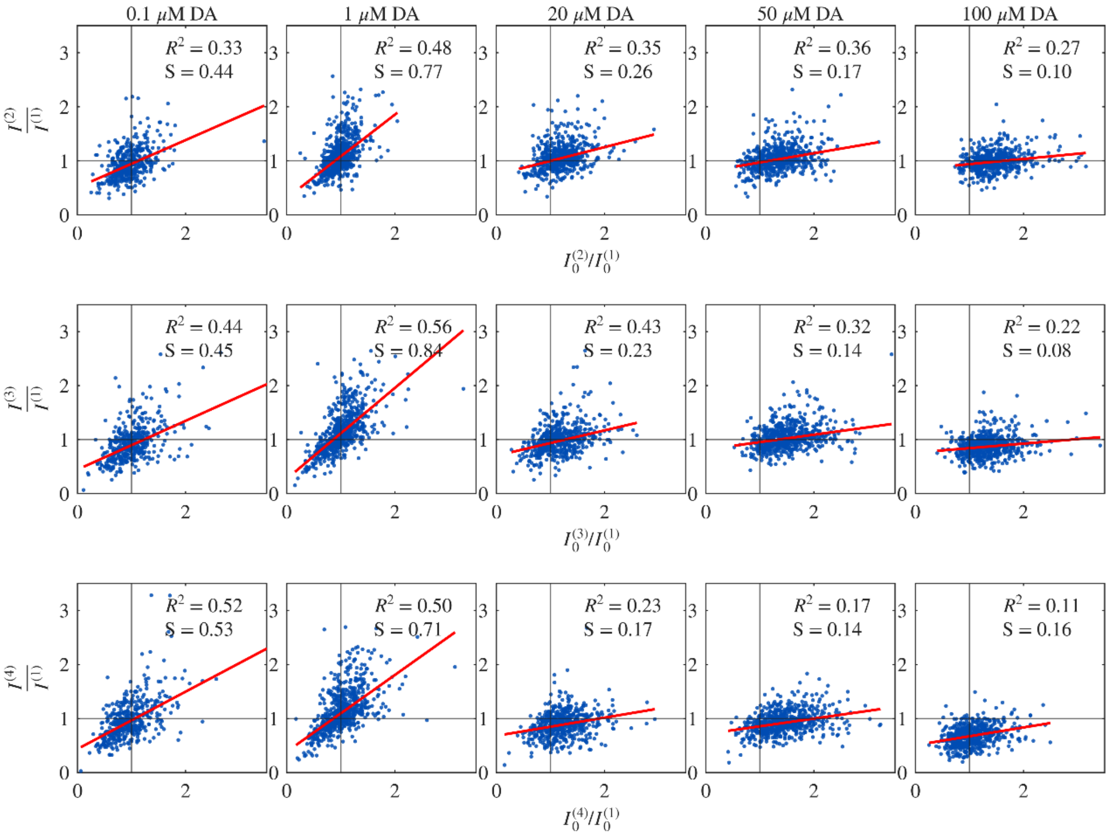


**Figure S13.** Comparison of the relative change in $I$ values to the relative change in $I_0$ values between the first and later cycles, for individual ROIs of DNA1-SWCNT and varying DA concentrations. Each dot represents an individual ROI. The black lines represent the $x = 1$ and $y = 1$ lines as guides to the eye. The coefficient of determination ($R^2$) of the data in relation to a linear fit, and the fit's slope ($S$) are presented, as well as the fitted curve (red).

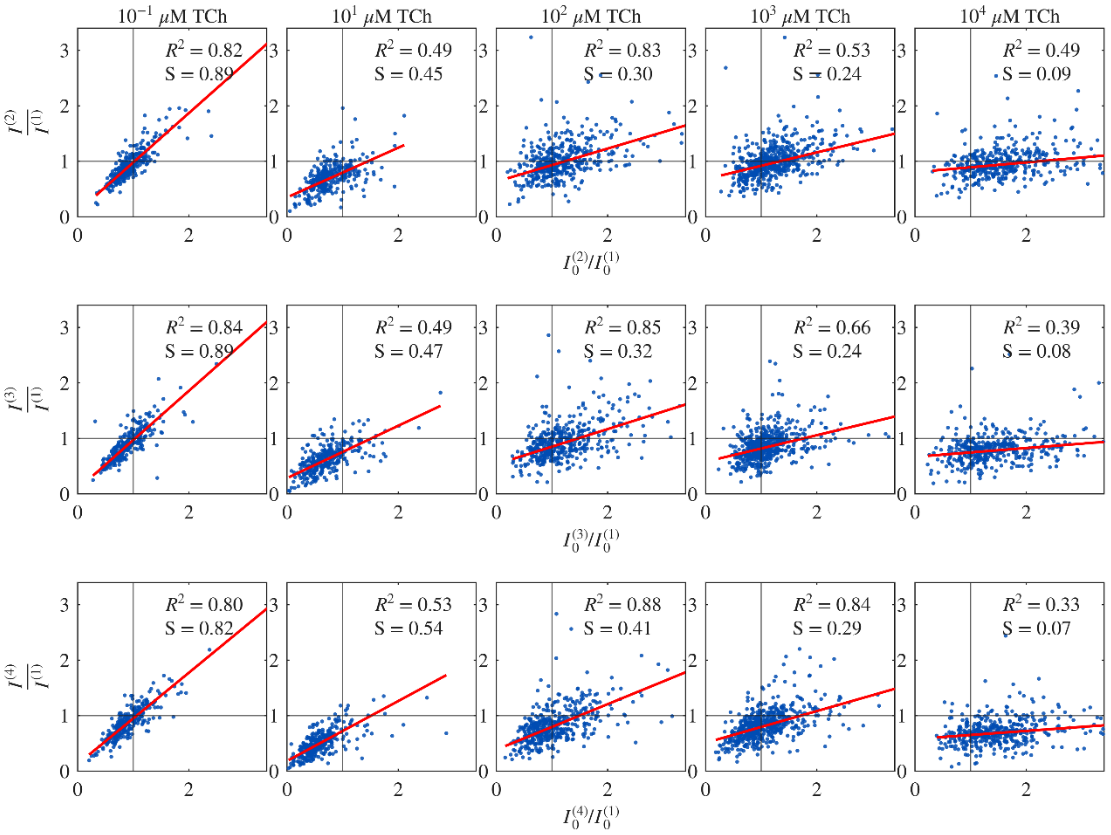


**Figure S14.** Comparison of the relative change in $I$ values to the relative change in $I_0$ values between the first and later cycles, for individual ROIs of DNA2-SWCNT and varying TCh concentrations. Each dot represents an individual ROI. The black lines represent the $x = 1$ and $y = 1$ lines as guides to the eye. The coefficient of determination ($R^2$) of the data in relation to a linear fit, and the fit's slope ($S$) are presented, as well as the fitted curve (red).

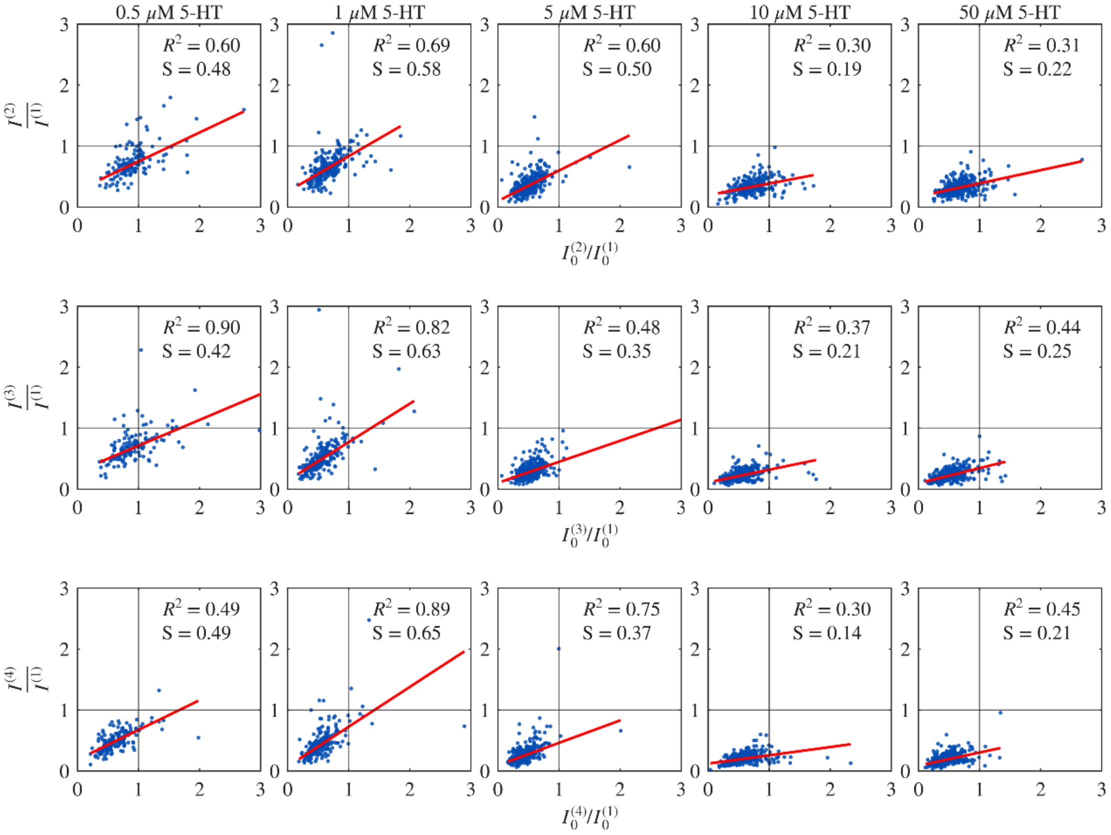


**Figure S15.** Comparison of the relative change in $I$ values to the relative change in $I_0$ values between the first and later cycles, for individual ROIs of DNA3-SWCNT and varying 5-HT concentrations. Each dot represents an individual ROI. The black lines represent the $x = 1$ and $y = 1$ lines as guides to the eye. The coefficient of determination ($R^2$) of the data in relation to a linear fit, and the fit's slope ($S$) are presented, as well as the fitted curve (red).

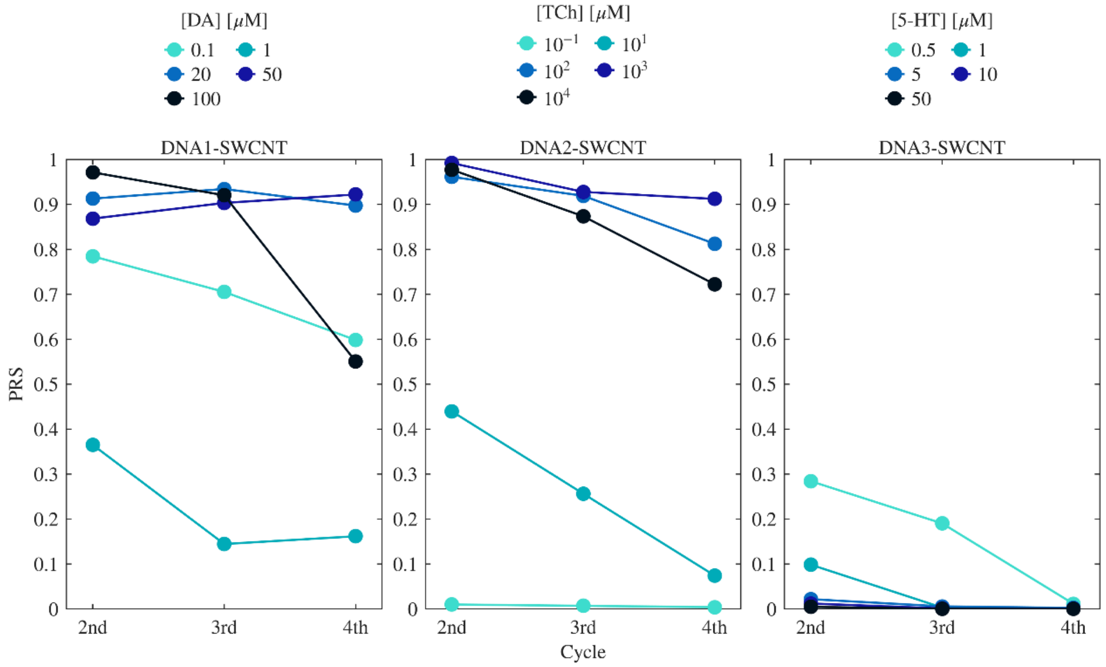


**Figure S16.** Reversibility Score of normalized fluorescence response, $(I - I_0)/I_0$, of the DNA-SWCNTs to varying analyte concentration across exposure cycles, using a fixed baseline $I_0^{(1)}$. From left to right: DNA1-SWCNTs with DA as an analyte, DNA2-SWCNT with TCh as an analyte, DNA3-SWCNT with 5-HT as an analyte.
